# Structural modelling of human complement FHR1 and two of its synthetic derivatives provides insight into their *in-vivo* functions

**DOI:** 10.1101/2022.11.24.517849

**Authors:** Natalia Ruiz-Molina, Juliana Parsons, Eva L. Decker, Ralf Reski

**Affiliations:** Plant Biotechnology, Faculty of Biology, University of Freiburg, Freiburg, Germany; Signalling Research Centres BIOSS and CIBSS, University of Freiburg, Freiburg, Germany

**Keywords:** AlphaFold, complement factor H-related 1, complement regulation, complement therapeutics, factor H, membrane attack complex

## Abstract

Human complement is the first line of defence against invading pathogens and is involved in tissue homeostasis. Complement-targeted therapies to treat several diseases caused by a dysregulated complement are highly desirable. Despite huge efforts invested in their development, only very few are currently available, and a deeper understanding of the numerous interactions and complement regulation mechanisms is indispensable. Two important complement regulators are human Factor H (FH) and Factor H-related protein 1 (FHR1). MFHR1 and MFHR13, two promising therapeutic candidates based on these regulators, combine the dimerization and C5-regulatory domains of FHR1 with the central C3-regulatory and cell surface-recognition domains of FH. Here, we used AlphaFold2 to model the structure of these two synthetic regulators. Moreover, we used AlphaFold-Multimer (AFM) to study possible interactions of C3 fragments and membrane attack complex (MAC) components C5, C7 and C9 in complex with FHR1, MFHR1, MFHR13 as well as the best-known MAC regulators vitronectin (Vn), clusterin and CD59, whose experimental structures remain undetermined. AFM successfully predicted the binding interfaces of FHR1 and the synthetic regulators with C3 fragments and suggested binding to C3. The models revealed structural differences in binding to these ligands through different interfaces. Additionally, AFM predictions of Vn, clusterin or CD59 with C7 or C9 agreed with previously published experimental results. Because the role of FHR1 as a MAC regulator has been controversial, we analysed possible interactions with C5, C7 and C9. AFM predicted interactions of FHR1 with proteins of the terminal complement complex (TCC) as indicated by experimental observations, and located the interfaces in FHR1_1-2_ and FHR_4-5_. According to AFM predictions, FHR1 might partially block the C3b binding site in C5, inhibiting C5 activation, and block C5b-7 complex formation and C9 polymerization, with similar mechanisms of action as clusterin and vitronectin. Here, we generate hypotheses and provide the basis for the design of rational approaches to understand the molecular mechanism of MAC inhibition, which will facilitate the development of further complement therapeutics.

## 1. INTRODUCTION

The human complement system is a fundamental part of innate immunity. It builds a surveillance network, which plays a key role in maintaining host homeostasis, contributing to the clearance of dead or modified host cells, and provides the first line of defence against invading pathogens.

Complement consists of more than 40 proteins in fluid-phase or associated to the surface of host cells. The complement system is activated in a cascading manner, and this can be initiated by three pathways, the classical (CP), lectin (LP), or alternative (AP) pathway, via proteolysis and/or conformational changes of the involved proteins. These signalling pathways converge at the activation of the very abundant complement component C3 and end up in inflammation, in opsonization of cells, tagging them for phagocytosis, and in membrane attack complex (MAC) formation, triggering lysis (reviewed by [1]).

While the CP and LP are activated upon recognition of invaders, the AP is permanently active in small amounts by spontaneous hydrolysis of the constitutively buried thioester bond in C3, leading to C3(H_2_O). Generation of C3(H_2_O) induces a conformational change of C3 and migration of the thioester-containing domain (TED), leading to a C3b-like molecule, which can bind Factor B (FB), but unlike C3b, it cannot attach to surfaces [2]. C3(H_2_O)-bound FB becomes activated by Factor D (FD) to generate a C3 convertase C3(H_2_O)Bb in the fluid phase, which cleaves C3 into C3a and C3b. C3a is an anaphylatoxin, which promotes cell recruitment and inflammation. C3b is a central component in the activation cascade and can bind to almost any surface in immediate proximity to the activation site and tag them for phagocytosis, a process known as opsonization, or it can form together with FB the surface-bound convertase C3bBb, establishing the amplifying loop of the AP, and extensively increasing the complement response (reviewed by [3]).

As a consequence of excess C3b, C5 is activated and the terminal pathway is initiated leading to the sequential non-enzymatic assembly of MAC, also known as terminal complement complex (TCC), which builds a lytic pore on membranes. C5 is activated either by cleavage to C5a, a potent anaphylatoxin, and C5b; or without proteolytical cleavage on highly dense opsonized cells, leading to a C5b-like activated C5 [4]. C5b can bind to C6, C7, C8, and C9 building a C5b6-9 arc and leading to polymerization of C9, MAC formation and cell lysis (reviewed by [5]).

As C3b cannot differentiate between foreign- and self-surfaces, cells that are not specifically protected, either by expressing regulators on their surfaces or recruiting them from plasma, can be marked and attacked by the complement system (reviewed by [6]). Therefore, the system is tightly controlled at different levels, in fluid phase and on host cells, to avoid damage of intact body cells, yet allowing efficient clearance of pathogens and damaged cells. Complement regulators typically display pattern recognition regions and binding sites for one or more complement components, and eventually other regulators. As the AP is responsible for 80% of the terminal complement activity [7], regulators of this pathway are of special interest.

Due to the ability of AP C3 convertases to amplify the complement response, their inactivation plays an important role in restricting complement activation to the correct target and appropriate degree. The main regulator in the fluid phase of the AP C3 convertases is Factor H (FH). FH consists of 20 globular domains, the short consensus repeats (SCRs). FH recognizes sialic acids and glycosaminoglycans on healthy host cells, binds them, mainly through SCRs 7 and 19-20 (FH_7_ and FH_19-20_), and targets its regulatory activity to protect body tissues. FH_1-4_ are able to bind to C3b and compete with FB, thus inhibiting convertase assembly. It can also displace the fragment Bb from the already formed AP C3 convertase, dissociating the complex, an activity known as decay acceleration (DAA). Moreover, FH displays a cofactor activity. When bound to C3b, FH acts as cofactor for the cleavage by Factor I of C3b to inactive C3b (iC3b), and subsequently to C3c and C3dg and finally the latter to C3d. The inactivated fragments are unable to bind FB. However, C3d, which contains the TED domain, can bind to FH through FH_19-20_ [8].

In contrast to the well characterized negative regulatory activity of FH, the regulation mechanism of the five FH-related proteins (FHRs) is still controversial. FHRs lack the C3b-regulatory region of FH, FH_1-4_, but they share high similarity to the C3d and surface recognition domain FH_19-20_.

FHR1 is the most abundant FHR, present in plasma in a similar concentration as FH [9], and of special interest. Due to the C-terminal homology to FH it was proposed to compete with FH for binding to C3b and host cell surfaces, displacing it and leading to local deregulation of complement activation [9]. However, it was recently shown that FHR1’s C-terminal domain, with only two substitutions compared to FH, is almost unable to recognize α2,3-linked sialic acids as host surface markers [10, 11]. Moreover, in contrast to FH, FHR1 was reported to bind C3, probably recruiting it to promote opsonization [11], and to bind to C5, C6, C7, C8 and C9, inhibiting MAC formation [12, 13]. Other inhibitors of MAC assembly include CD59, vitronectin and clusterin. FHR1 displays a dimerization domain in the N-terminal domains (FHR1_1-2_), and it can circulate as homodimer or heterodimer together with FHR2 or FHR5 [9].

Although one of the main functions of the complement system is to protect the body from pathogens, many microorganisms have developed a diverse range of strategies to evade complement attack by, e.g., recruiting soluble complement regulators as a protective shield (reviewed by [1]). On the other hand, over-activation of complement in response to viral infections such as hepatitis C, dengue and coronavirus has been associated with viral pathogenesis [14–17]. Moreover, mutations in complement-associated proteins and deregulation of the activation cascade are associated with a long list of diseases such as age-related macular degeneration, atypical haemolytic uremic syndrome, C3 glomerulopathies, paroxysmal nocturnal hemoglobinuria, and systemic lupus erythematosus. Furthermore, dysregulation of complement activation can contribute to autoimmune and inflammatory diseases [18].

Despite vast efforts, only very few complement-related drugs have reached final regulatory approval. Therefore, there is growing interest in recognizing interactions between complement effectors and regulators, to better understand the role of complement in disease. This knowledge can facilitate the rational development of therapeutics to restore homeostasis in different diseases associated with complement dysregulation.

FH and FHRs could be targets for therapeutic approaches (reviewed by [19]). However, due to the structural complexity of FH, its recombinant production is far from trivial [20, 21]. Therefore, smaller proteins including only the main active domains of FH have been developed [22–24].

More recently, two multitarget fusion proteins, MFHR1 (FHR1_1-2_:FH_1-4_:FH_19-20_) and MFHR13 (FHR1_1-2_:FH_1-4_:FH_13_:FH_19-20_), were developed, which combine the dimerization and C5- regulatory domains of FHR1 with the C3-regulatory and cell surface recognition domains of FH to regulate the activation of the complement in the proximal and the terminal pathway, and target the regulation to host surfaces. These fusion proteins exhibited a superior overall regulatory activity *in vitro* compared to FH [13,25,26].

The artificial intelligence AlphaFold2 (AF2) has recently revolutionized structural biology due to unprecedented success and accuracy in protein fold prediction [27]. AF2 is a neural network that uses physical, geometric, and evolutionary constraints. AF2 uses co-evolution information from multiple sequence alignments (MSAs) and was furthered trained to predict protein-protein interactions (PPI) by an algorithm called AlphaFold-Multimer (AFM) [28]. While proteins can adopt different states under specific conditions, only one single state is predicted by AF2/AFM, which is one of the major limitations.

ColabFold was developed based on AF2 and AFM with some modifications, and enables faster prediction in local machines without compromising accuracy [29]. Benchmarking of AFM showed significantly higher accuracy compared with global docking approaches, with success rates varying between 51% and 72% [28,30,31]. However, there is still a low rate of success for antibody-antigen complexes, probably due to lack of coevolution signals in MSAs (11%) [31]. This is also the case for individual proteins without enough coevolutionary relationships such as metagenomic protein sequences and viral proteins. Recently, a new algorithm was created to overcome these limitations [32].

Here, we studied the structures and interactions of MFHR1 and MFHR13 with different complement components to correlate structural features with previous experimental observations [13]. Furthermore, many structures of MAC regulators and their complexes remain experimentally undetermined. Therefore, we included AFM predictions of the most important MAC regulators, including FHR1, analysed the interactions with MAC components C5, C7 and C9, and generated new hypotheses about the mechanisms of MAC inhibition taking into account previously published experimental results.

## 2. RESULTS AND DISCUSSION

### 2.1 Comparison of MFHR1 and MFHR13 structure models

To further characterize the synthetic complement regulators MFHR1 and MFHR13, we analysed their structure. The rational design of MFHR13 originally involved the modelling of the protein structure using Modeler 9.19 [13]. Here, we used ColabFold based on the AlphaFold2 (AF2) algorithm and AlphaFold-Multimer (AFM) to model both chimeric proteins and evaluate their ability to dimerize.

The AF2 algorithm generated five top-ranked models for MFHR1 and MFHR13. These top-ranked models have a similar conformation and orientation of domains SCRs 19-20 of FH (FH_19-20_) for MFHR1 in contrast to MFHR13 (Figure 1 A, B, Supplementary Figure 1 A). MFHR13 models differ in the orientation of FH_13_, FH_19_, and FH_20_, probably due to the flexibility of the linkers between SCRs in the C-terminus of the protein. The linker between FH_13_ and FH_19_ in MFHR13, equivalent to the linker between FH_13_ and FH_14_ (13-14 linker) in FH (SMAQIQL), was always predicted with very low confidence (pLDDT< 50), contrary to the linker between FH_4_ and FH_19_ in MFHR1, which is the natural linker between SCR 18 and 19 (18-19 linker) in FH (EDSTGK). Very low confidence regions predicted by AF2 have been correlated with intrinsically disordered regions (IDRs) [33], which can acquire several conformational ensembles and evolve faster than ordered regions. Furthermore, 13-14 linker in FH is the most variable across species; for instance, the sequences are NEEAKIQL and KEEVLNS in cows and pigs, respectively [34]. Likewise, FH_13_ itself is variable in sequence between orthologues, and the structure differs from the rest of the SCRs. Maximum eight β-strands (named A – H) have been recognized in SCR structures. FH_13_ is the most spherical SCR and lacks β-strand H, along with a three-residue deletion in the flexible loop between β-strands D and E [34]. This loop and the β- strand H are important to stabilize the interface with a consecutive SCR. These unique features indicate that FH_13_ can contribute to the structural flexibility of MFHR13, in agreement with the flexibility predicted by AF2 for the interface 13-19 in this fusion protein.

**Figure 1.**
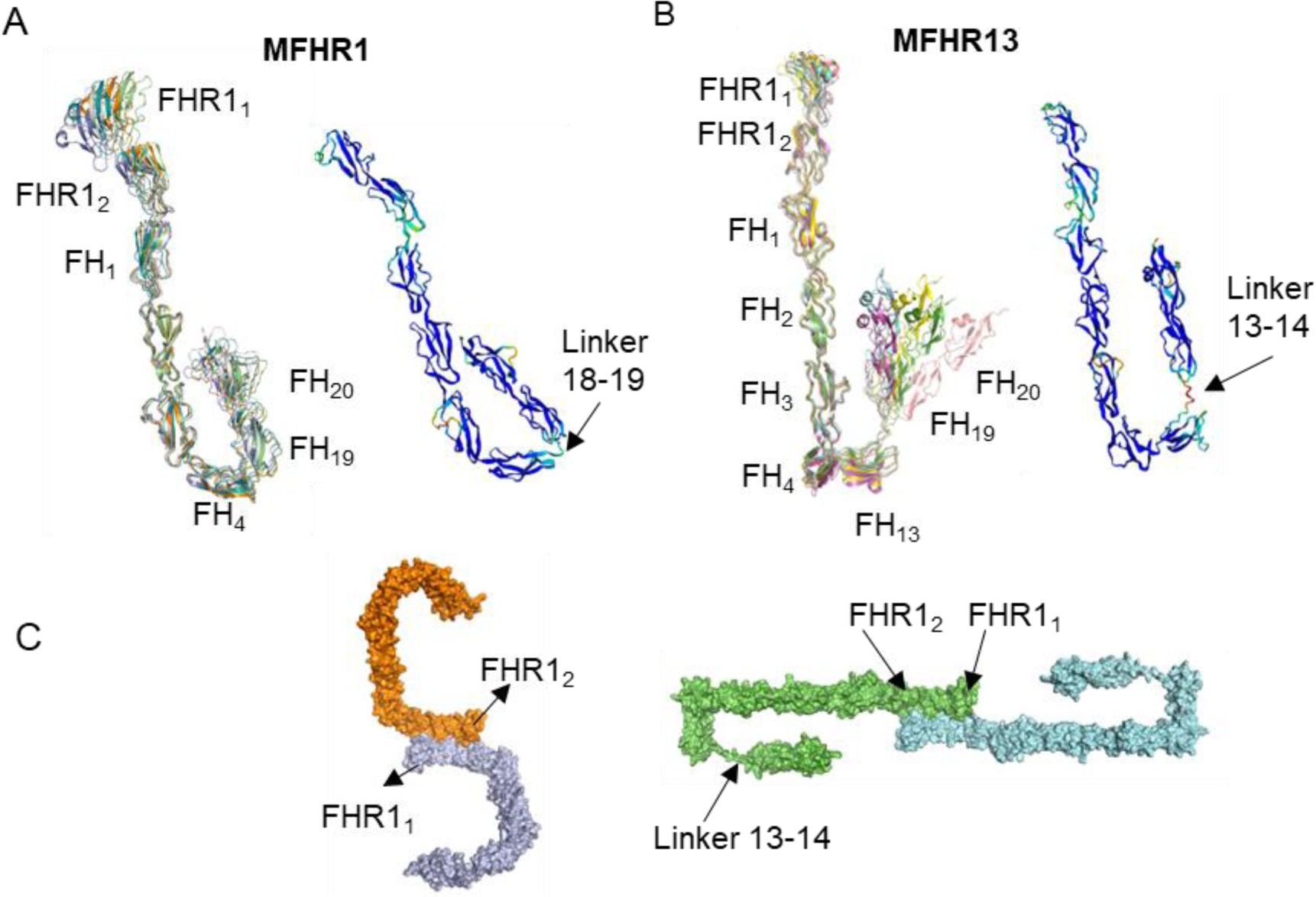
Structure models of the synthetic complement regulators MFHR1 and MFHR13 predicted by AlphaFold2 (AF2). Structure of MFHR1 top-ranked models (**A**) and MFHR13 (**B**). Superimposition of the 5 top-ranked models are shown on the left, while the best model is displayed on the right with the pLDDT score (0-100) by colours (>90 in blue, high confidence, and <50 in red, low confidence). **C.** Prediction of MFHR1 on the left and MFHR13 on the right dimerization interface by AlphaFold-Multimer (AFM). The linker between FHR1 and FH domains is also predicted with different possible conformations.

The quality of the top-ranked model was further evaluated with QMEANDisCo and PROCHECK (Supplementary Figure 1 B, C). QMEANDisCo includes a new distance constraint score based on pairwise distances in the model and constraints obtained from PDB structure homologues to the model. The low score of FH_13_ and the linker might be due to a few close homologues with available experimental structures, which decrease the accuracy of QMEANDisCo [35]. The superimposition of PDB structures (PDB 2wii, 2kms, 2xqw) and AF2 prediction with a backbone root-mean-square deviation (RMSD) of 1.37, 0.93, and 1.2 Å for FH_1-4_, FH_13_, and FH_19-20_, respectively, indicates that they are quite similar to the experimental structures (Supplementary Figure 2 A). Furthermore, AFM correctly predicted the dimerization interface in FHR1_1-2_ on both synthetic regulators compared with an experimental structure of FHR1_1-2_ (PDB 3zd2) (Figure 1C), although the prediction quality was not highly confident (Supplementary Figure 2 B).

### 2.2 Binding of MFHR13, MFHR1 to C3 and C3 cleavage fragments

C3 consists of two chains (α and β), containing 13 domains; eight of them with a fibronectin-type 3-like core fold, called macroglobulin (MG) domains. A series of proteolytic reactions results in several C3 fragments with different functional properties. These fragments are derived from the α-chain while the β-chain remains intact. MG1-MG5 and part of MG6 form the β-chain with a linker domain. The α-chain is formed by an anaphylatoxin domain, part of MG6, along with MG7, MG8, a thioester-containing domain (TED), complement C1r/C1s, Uegf, Bmp1 domain (CUB), and the C-terminal domain, called C345C [36]. The anaphylatoxin domain forms C3a when C3 is cleaved by C3 convertases, and the TED domain forms C3d when C3b is cleaved for inactivation, and allow covalent attachment of C3b and C3d to cell surfaces (opsonization).

It is well known that FH can bind C3b through FH_1-4_ or FH_19-20_ and regulate complement activation [37]. In order to delineate a probable mechanism for the synthetic regulators MFHR1 and MFHR13, we evaluated their binding to C3b using AFM and compared it to the experimental structures of FH_1-4_/C3b (PDB 2wii), FH_19-20_/C3d (PDB 2xqw), and FH_18-20_/C3d (PDB 3sw0).

AFM predicted the interaction between MFHR13 and C3b with interfaces located in FH_1-4_ and FH_19_ (Figure 2 A) in agreement with experimental structures of FH fragments [8, 38]. According to the model, both sites might be used simultaneously to bind one C3b molecule, while these binding sites are considered independent in FH [23]. The ability to bind simultaneously to one C3b molecule can have an impact on the functional activity of the protein as a potential therapeutic agent, which will be further discussed. The experimental structure of FH_1-4_/C3b and the model of MFHR13/C3b are highly similar with an RMSD of 1.253 Å, but the orientation of FH_4_ is slightly different (Figure 2 A, zoomed panel). The linkers in FH_1-4_ have some degree of inherent flexibility and it has been shown that a kink between FH_3_ and FH_4_ occurs when they bind C3b [37]. An additional C3b binding site in FH was suggested to be located in FH_13-15_, which binds with a weak affinity [39], however none of the analysed models predicted a binding site localized in FH_13_ in MFHR13.

**Figure 2.**
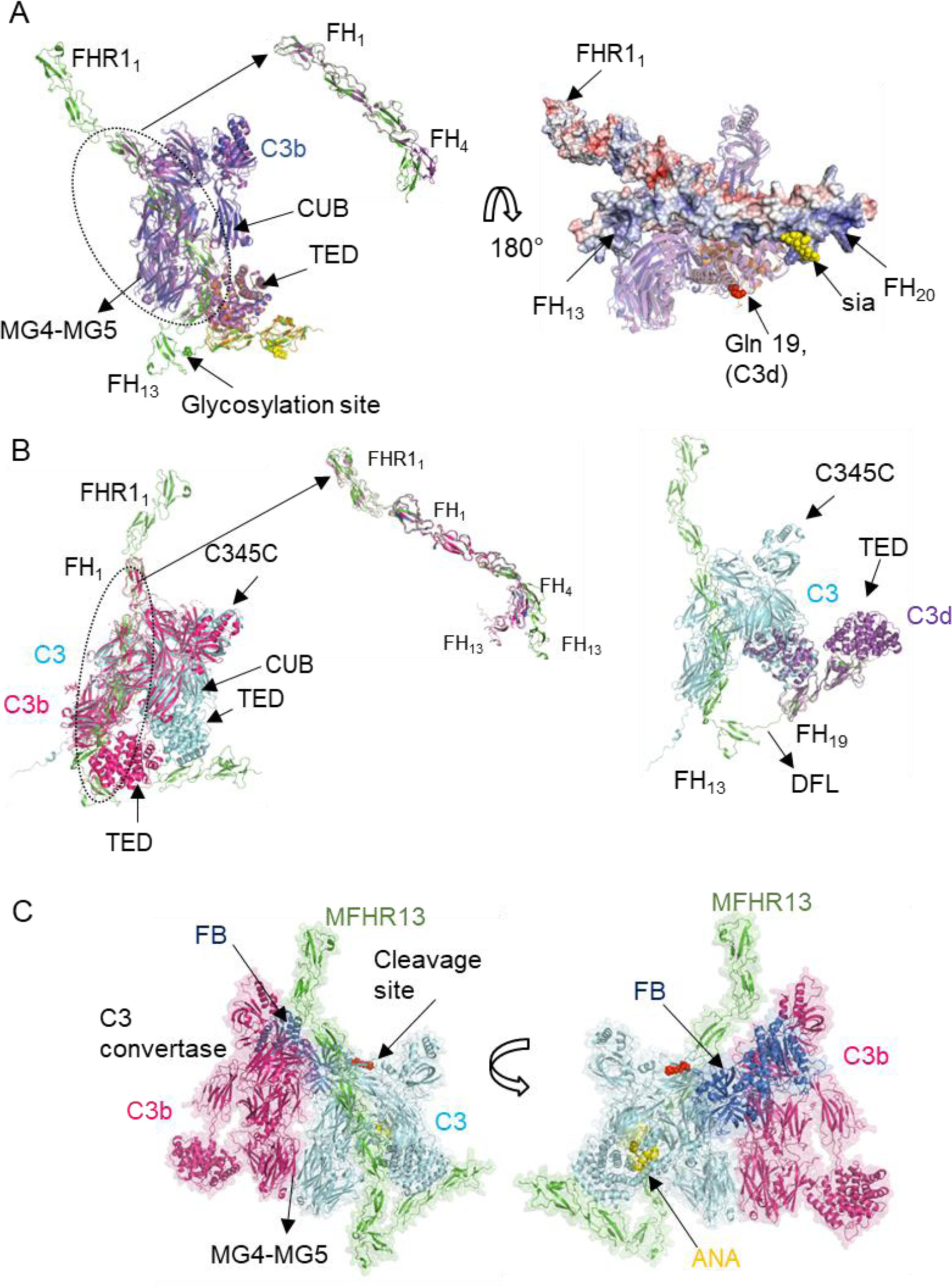
Interaction of MFHR13 with C3 and C3b predicted by AlphaFold-Multimer. **A.** The model of the complex MFHR13/C3b predicted by AFM agrees with experimental structures of FH fragments in complex with C3b. Superimposition of the model MFHR13 (green) interacting with C3b (blue) with experimental structure of FH_1-4_/C3b (magenta) (PDB 2wii, RMSD = 1.253 Å), FH_19-20_/C3d (orange) (PDB 2xqw, RMSD = 0.928 Å) and sialic acid (yellow) (PDB 4ont), RMSD = 1.253 Å). The thioester-containing domain (TED) and complement C1r/C1s, Uegf, Bmp1 domain (CUB) are indicated by arrows. Zoom in of the superimposition of FH_1-4_ of the model and the experimental structure is shown. On the right are shown the linkage of C3b (or C3d) to biological surfaces around Gln19 (red spheres) and MFHR13 as a surface with the electrostatic surface potential (electropositive residues in blue and electronegative in red). **B**. MFHR13 interacts with C3 through FH_1-3_ and FH_19_ domains. On the left, MFHR13 and C3 are shown in green and cyan, respectively, and the model is superimposed with C3b in complex with FH_1-4_ (PDB 2wii, in magenta), where differences in C3 and C3b conformations are observed.

We compared the models of MFHR1 and MFHR13 in complexes with C3b. Although AFM predicted the correct interfaces for C3b binding on FH_1-4_ and FH_19-20_, the differences in flexibility of the 18-19 linker in MFHR1, and the 13-14 linker in MFHR13, respectively, determine if the binding to one C3 or C3b molecule occurs independently or simultaneously via both interfaces.

The model of MFHR1 in complex with C3b showed a different orientation of FH_4_ and FH_19_ compared with FH_18_ and FH_19_ in the crystal structure FH_18-20_ (PDB 3sw0) (Supplementary Figure 3 A). Probably, the conformation of FH_18-19_ in the experimental structure (PDB 3sw0) is just one of several possible conformations, since the 18-19 linker is significantly more flexible than the19-20 linker [40]. Moreover, a mini-FH version with an optimized artificial linker (12 glycines) outperformed the original version with the natural linker 18-19 in an overall alternative pathway regulation assay, which was attributed to the inability of the original molecule to bind simultaneously to one C3b molecule using both interfaces [41]. We conclude that in MFHR1 the simultaneous interaction of one C3b molecule with both interfaces (FH_1-4_ and FH_19_) is theoretically possible, although the short length of the 18-19 linker can impose some constraints, which could be overcome by the use of an optimized linker. Although there was no experimental difference between MFHR1 and MFHR13 in C3b binding *in vitro* by an ELISA-based method [13], other approaches like Surface Plasma Resonance (SPR) or Bilayer interferometry (BLI) should be evaluated to determine the differences in association and dissociation rate to C3 and its fragments.

It is important to highlight that AFM predicted a simultaneous binding between C3b and MFHR1 or MFHR13 through both interfaces only by using templates. The amount of recycles plays an important role as it was previously described [31] and up to 15 or 8 recycles were needed for MFHR1 and MFHR13, respectively, otherwise only one of the interfaces were predicted. The Inter-PAE for the models are shown in Supplementary Figure 3 B.

According to the model, when MFHR13 binds to surface-bound C3d or C3b, the electropositive patch between FH_13_ and FH_20_ is oriented towards the surface to bind GAGs or sialic acid, leaving the active domains to regulate complement on host cell surfaces (Figure 2 A, right). We observed a higher binding to heparin which is correlated with the presence of FH_13_ in MFHR13 compared with MFHR1 [13].

While FH cannot bind C3, it was recently shown that FHR1 binds C3, C3b, iC3b and C3d [11]. Therefore, we included C3 as a potential ligand of MFHR1 and MFHR13. AFM without templates predicted an interaction of C3 with interfaces located in FH_1-3_ and FH_19_ of MFHR13 and MFHR1 (Figure 2 B, Supplementary Figure 3 C, D). The interaction between FH_4_ and the TED domain, which occurs in the complex FH_1-4_/C3b (PDB 2wii) [42], might be prevented in MFHR13 due to conformational differences between C3b and C3, specifically in the C345C and TED domains (Figure 2 B). C3-binding sites in FH might be cryptic and thus FH cannot bind C3. However, the fragment FH_1-6_ binds C3, but only with 5-fold weaker affinity compared to FH/C3b [39], which is in agreement with the AFM prediction. Thus, we propose that in contrast to FH, but similar to FHR1 or FH_1-6_, MFHR13 and MFHR1 might bind to C3 due to their extended conformation in fluid phase. As opposed to MFHR13, it is unlikely that MFHR1 can interact with one C3 molecule simultaneously through the interfaces predicted in FH_19-20_ and FH_1-3_, since the 18-19 linker would need to acquire a complete extended conformation (Figure 2 B, Supplementary Figure 3 C).

Although AF2 has an outstanding ability to predict global protein structures, the accuracy decreases significantly for multi-domain proteins compared to individual domains [43, 44].

Therefore, inter-domain orientation in models of MFHR1, MFHR13, FH and FHR1 should be interpreted with caution. MFHR13 structure models always showed a similar orientation of FH_3_, FH_4_ and FH_13_. However, in complex with C3 ligands, the orientation of FH_4_ with respect to FH_3_ changed slightly, along with the flexible 13-14 linker which acquired an extended conformation to allow the simultaneous binding to one C3 molecule through both interfaces (FH_1-4_ and FH_19_) (Figure 2 A, B). Although an extended linker conformation may not be reliable, this linker is likely to be a disordered flexible linker (DFL) according to the pLDDT as mentioned above.

Despite other approaches such as IUPred2A, DISOPRED3, fMoRFpred, DFLpred and TransDFL failed to predict this linker as a disordered region (Supplementary Figure 3 E), a previous study showed that AF2 exceeded the performance of 11 disorder region predictors on the DisProt-PDB dataset [45].

RMSD is 2.583 Å for the whole complex superimposed with PDB 2wii, and 1.376 Å for FH_1-4_, respectively. On the right superimposition of MFHR13/C3 model (green/cyan) with experimental structure FH_19-20_/C3d (PDB 2qxw, in purple) is shown. RMSD is 0.601 Å for C3 TED domain and C3d. The disordered flexible linker (DFL) is shown. **C.** C3 convertase (C3bBb) in complex with C3/MFHR13 (enzyme-substrate complex). Superimposition of C3 in MFHR13/C3 model with MG4-MG5 domains of C3b molecule in C3 convertase (PDB 2win, C3b in magenta, Bb fragment in dark blue). MFHR13 is shown in green and C3 in light blue with the anaphylatoxin domain (ANA) in yellow and the cleavage site as red spheres.

The cofactor activity is essential for complement regulation since C3b is inactivated to prevent C3 depletion, additional C3 convertases formation and C3b deposition on cell surfaces [46]. FH and the synthetic regulators MFHR1 and MFHR13 act as cofactor for factor I (FI), a serine protease, which circulates in an inactive form. The interactions between FI, C3b and FH are fundamental to triggering the remodelling of the active site [47]. Different mini-FH versions exhibited lower cofactor activity in fluid-phase than full-length FH [22, 48], as is the case for MFHR1 and MFHR13 [13,25,26]. According to AFM models we speculate that a high entropy in MFHR13 due to the 13-14 linker might destabilize the interaction between C3b and FH_4_ when FH_19_ is interacting simultaneously with the TED domain of C3b. Although FH_4_ in FH is not directly interacting with FI as FH_1-3_ is, FH_4_ interaction with C3b might hold the TED domain position while the CUB domain is cleaved [37]. In contrast, this conformation of FH_4_ would not affect decay acceleration activity (DAA) because the interaction of FH_1-2_ with C3b can displace the Bb protease fragment (by electrostatic repulsion), while the other interactions of FH_3-4_ and even the FH_19_ can support the binding to C3b to improve the DAA. This might explain why MFHR13 was better than MFHR1 in DAA [13], since the simultaneous binding to C3b is more feasible than for MFHR1.

The ability of MFHR1 and MFHR13 to bind C3 as predicted by AFM might have implications for complement regulation that should be carefully considered. Surface-bound FHR1 can bind C3b, allowing convertases assembly by displacing FH, and it can also bind to C3, increasing its local concentration, generating more C3b and in turn triggering complement activation instead of negatively regulating it [11, 49]. MFHR13 can bind C3 not only through the TED domain as FHR1 does. Therefore, to check if MFHR13 binding to C3 avoids its cleavage by C3 convertases, we created a hypothetical model of C3 convertase interacting with C3 as described by [50].

MFHR13 bound to C3 would not interfere with the interaction of C3 and C3 convertases whose interface is located in MG4-MG5 domains. Thus, it would not prevent cleaving additional C3 into C3b, similar to what was shown for FHR1 variants [10] (Figure 2 C). Once C3b is generated, MFHR13 can act as cofactor for FI to inactivate C3b to iC3b, thus counteracting complement activation, which is not possible for FHR1. Although it applies also for MFHR1, the affinity of both synthetic regulators to C3 should be determined experimentally. Even if both proteins perform well as complement regulators *in vitro*, these characteristics should be considered when evaluating different performances *in vivo*.

Previously, we also analysed the binding to C3d, an inactivated C3b fragment, which is important for MFHR1 and MFHR13 to target the therapy to regions with ongoing complement activation [13]. It is well known that FHR1 and FH bind C3d through FHR_4-5_ [51] and FH_19-20_ [8], respectively. AFM without templates correctly predicted the interacting interfaces between C3d and those domains in FHR1, MFHR1, or MFHR13 with a good confidence score (Figure 3 A, B, C), although always lower for the synthetic regulators (Supplementary Figure 4). Additionally, AFM predicted an interface located in FHR1_1-2_, which is also present in the synthetic regulators (Figure 3 A, B, C, D). However, the corresponding interface on C3d would be blocked since it is linked to biological surfaces through a region around Gln19. Therefore, it might not be physiologically relevant and it is predicted with a lower score compared to FHR1_4-5_ (Supplementary Figure 4). Furthermore, the extended conformation of 12-13 linker in MFHR13 is not reliable, since it is a not flexible linker [34], therefore the protein might not interact simultaneously with one C3d molecule using both interfaces. The interface in FH_19-20_ coincides with the experimental structure (PDB 2xqw), and more residues in the interface FH_19-20_ are interacting in AFM models as in the experimental structure, which derived from the position of the lateral chains (Figure 3 E).

**Figure 3.**
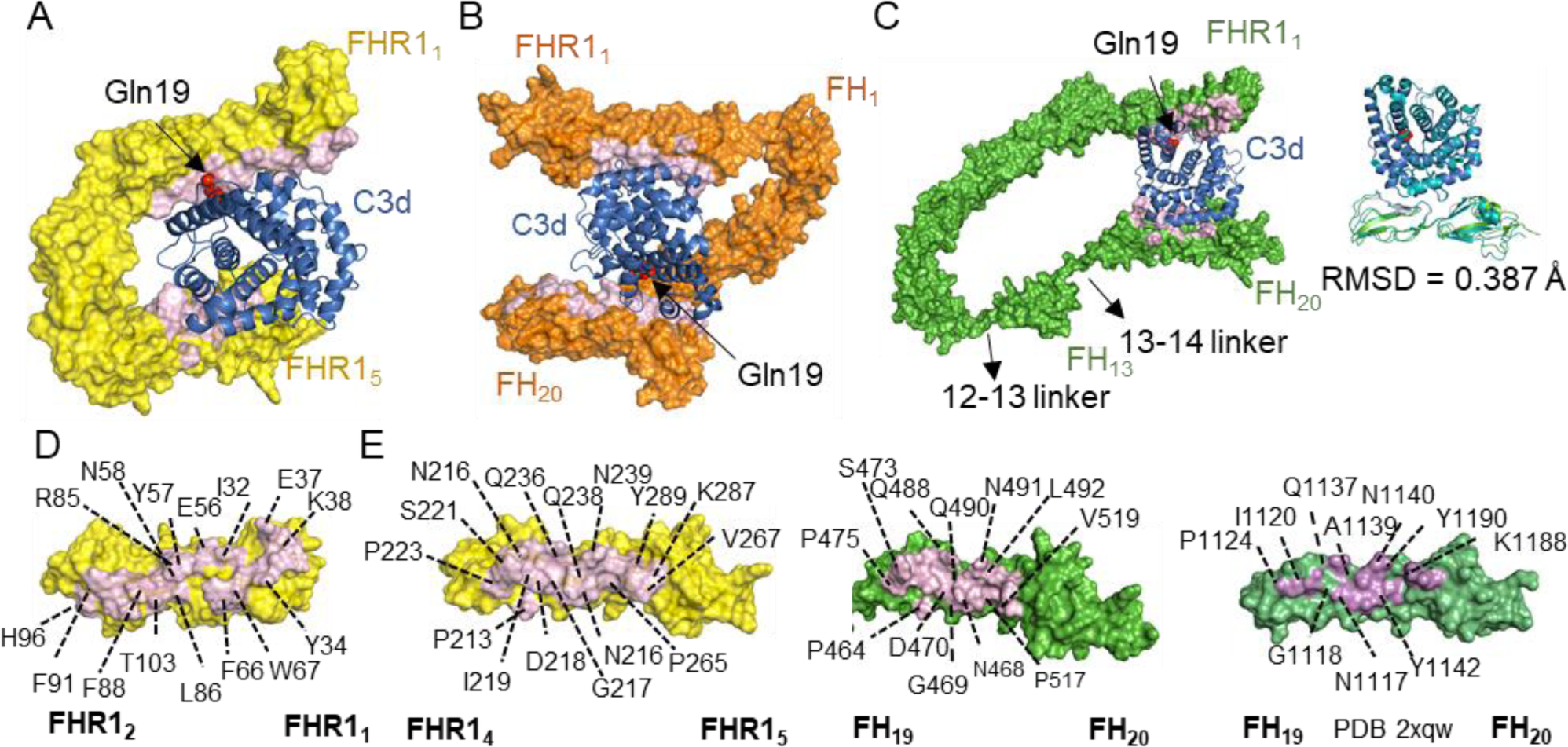
Interactions of FHR1, MFHR1 and MFHR13 with C3d predicted by AlphaFold-Multimer. **A.** FHR1 interacts with C3d through FHR1_4-5_ with high confidence score while a second interface in FHR1_1-2_ was predicted overlapping the C3d region linked to biological surfaces around Gln19 (shown as red spheres in all structures). MFHR1 **(B)** and MFHR13 (**C**) interact with C3d through FH_19-20_ with a high confidence score. The same interface in FHR1_1-2_ was identified for these proteins **D**. Binding interface (in pink) between FHR1_1-2_ and C3d with some residues labelled in FHR1. **E.** Binding interfaces between FHR_4-5_ (on left) and FH_19-20_ with C3d predicted by AFM without templates (centre) compared with the experimental structure PDB 2xqw (on right). Some residues in the interface are labelled in FHR1.

### 2.3 Interactions between FHR1 and terminal complement components

The terminal pathway of complement activation leads to MAC formation, which builds a pore on the cell membrane triggering lysis of pathogens. MAC consists of C5b, C6, C7, the heterotrimeric C8 (C8αβγ) and around 18 C9 molecules [5]. The ability of FHR1 to bind C5 has been a matter of controversy [9, 12]. Recently, we observed binding of FHR1 as well as the synthetic proteins MFHR1 and MFHR13 to C5 and other components of the MAC. Moreover, in molar excess they were able to inhibit activation of the terminal pathway and cell lysis in a haemolytic test [13]. This indicates a role of FHR1 in regulating MAC formation.

AFM successfully predicted the interfaces of MFHR1, MFHR13 and FHR1 with C3, C3b and C3d. Further, we modelled their complexes with C5, C7 and C9 to evaluate whether the structures can provide insights into the molecular mechanism of MAC inhibition. Vitronectin (Vn) and clusterin are important soluble regulators of the MAC, and CD59 is the only regulator anchored to the membrane and is found on almost all tissues. How the MAC regulators work is still the subject of several studies but no crystal structures are available. Therefore, we included models of these regulators in our analysis.

The models were run using amber algorithm to place the side chains, with 3, 6, 12 and 20 recycles with and without templates and the 5 top ranked models predicted by AFM were analysed for each run.

#### 2.3.1 Interactions between FHR1 and C5

Similar to C3, C5 consists of 11 domains, 8 MG domains, the anaphylatoxin C5a, CUB domain, TED domain (C5d), and the carboxy-terminal C345C domain, which is flexibly attached to the C5 core (Figure 4). After activation, C5b and C5b-like are not stable (the half-life of C5b is around 2 min) and C6 should quickly bind to form a stable complex, C5b6, which is the first step in MAC formation [52]. The C5d domain interacts extensively with C6 to form C5b6 and, unlike C3b or C3d, C5b cannot covalently attach to cell surfaces through the TED domain, since it does not contain an internal thioester bond but a structural homologue of the TED domain of C3 [53].

**Figure 4.**
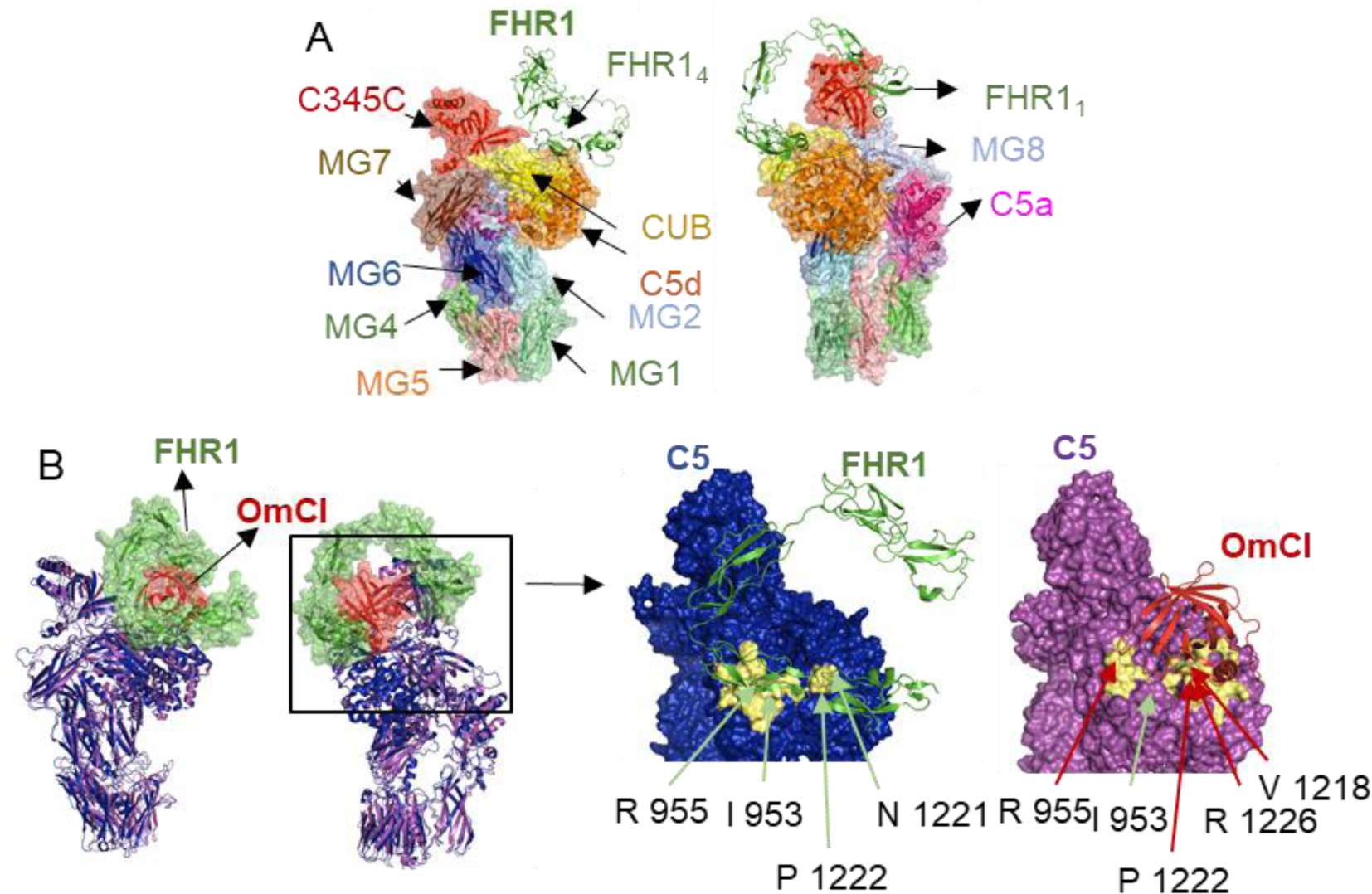
Interaction of C5 and FHR1 predicted by AlphaFold-Multimer. **A**. Model of the FHR1/C5 complex predicted a binding interface in CUB and C5d domains. The colours of C5 domains; shown as surface, match the colour of their legends and FHR1 is shown as cartoon (green). **B**. Model of the FHR1/C5 complex indicates a binding interface in the CUB and C5d domains, similar to complement inhibitor OmCI. Superimposition of the model FHR1/C5 predicted by AFM (C5 cartoon in blue, FHR1 surface in green) and C5 in complex with OmCI (PDB 6rqj) (C5 cartoon in purple, OmCI surface in orange), (RMSD = 0.742 Å). Binding interfaces of FHR1/C5 and OmcI/C5 in pale yellow with key residues indicated by green or red arrows, respectively, are shown on the right.

The top ranked AFM model predicts FHR1 interaction with C5 through interfaces located in FHR1_4-5_, where mainly 7 residues are involved forming 10 hydrogen bonds (Figure 4 A, Supplementary Figure 5). The top key residues predicted in the interface were Phe 222, Tyr 289, Gln 238, Gln 236, Thr 220, Pro 223 in FHR1 and Arg 955, Ile 953, Pro 1222, Phe 1352, Asn 1221, Thr 952 in C5. According to the AFM model, FHR1 would not bind near convertase- binding sites in C5 MG4-MG5 domains. Therefore, FHR1 would not interfere with convertase- binding by steric blocking. However, the model shows a similar binding interface as the complement inhibitor OmCI (Coversin, Nomacopan), isolated from the tick *Ornithodoros moubata*, which is currently under clinical trials to treat several complement-associated diseases [54]. This 16-kDa protein binds a region of C5 CUB and C5d and there is one amino acid involved in the C345C domain [52] (Figure 4 B). C5d-CUB-MG8 domains undergo a conformational rearrangement after proteolytic cleavage of C5 to C5b or non-proteolytic C5 activation, which is critical for C6 binding [4, 53]. It has been proposed that OmCI generates an allosteric modulation of convertases, reducing the affinity or blocking the binding of these to C5 [55]. Furthermore, the interaction with one residue in C345C seems to be critical for OmCI’s role as complement inhibitor [52]. Interestingly, in our models there is no FHR1 interaction with this domain.

According to the AFM model, FHR1 binds CUB and C5d domains with a smaller interface compared to OmCI (Figure 4 B), which binds to C5 with a high affinity constant (K_d_) in the low nanomolar range [56]. MFHR13, on the other hand, binds C5 with a K_d_ in the low micromolar range [13], which might limit its activity on the C5 level to elevated local concentrations of activated C5.

The molecular mechanism of proteolytic and non-proteolytic C5 activation is not well understood. C5 activation occurs on cell surfaces with C3b deposition, where interactions between C5 and C3b are critical. According to [4], C3 convertases cleave C5 in presence of additional C3b molecules. Experimental structures of C5 in complex with C3b are not available, but the structure of C5 in complex with cobra venom factor (CVF), a C3b homologue, suggests that C5 interacts directly with C3b, also as part of C3 convertases, through MG4-MG5 domains, similar to the C3 convertase in complex with C3 [57]. Other suggested binding sites for C3b are located in the C5 CUB domain [4]. Therefore, according to the AFM predicted model, we propose that binding of FHR1 to C5 might partially block C3b binding, inhibiting in turn C5 activation, which should be experimentally validated.

#### 2.3.2 Interactions between FHR1 and C7

C7 initiates the MAC assembly by binding to C5b6, forming C5b-7 which can attach to membranes and can recruit C8. C7 is a limiting factor in MAC assembly as it is the only TCC protein which is not mainly produced by hepatocytes [58] and C8 is an inhibitor of the MAC assembly itself because it can prevent the insertion into the membrane if it binds to C5b-7 before the insertion occurs [59].

C7 consists of nine domains, two thrombospondin-type domains (TSP type 1), a lipoprotein receptor class A domain (LDLRA), a membrane attack complex-perforin domain (MACPF) (also present in C6, C8 and C9), an epidermal growth factor (EGF) domain, and at the C-terminus of C7 there are two short-consensus repeats (SCRs) and two factor I-like membrane attack complex (FIM) domains which interact with C5b in the assembled MAC [60, 61]. MACPF is responsible for the β-barrel pore formation containing a helix-turn-helix (CH3) motif and two amphipathic transmembrane β-hairpins (TMH1, TMH2), while the rest act as auxiliary domains interacting with C6 or C8. Top-ranked models predicted by AFM show three potential binding interfaces between FHR1 and C7. However, the interface predicted with the highest score confidence is located in SCR1 and SCR2 of C7 comprising 20 residues (around 604-681), and 11 and 8 residues in FHR1_4_ and FHR1_5_, respectively (Figure 5 A, B, Supplementary Figure 6 A) forming 10 strong hydrogen bonds. Key residues in the interface include Arg 314, Thr 220, Phe 222, Asp 218 and Arg 291 in FHR1 and Ser 675, Ile 606, Gln 627, Ile 629, Ser 676 in C7.

**Figure 5.**
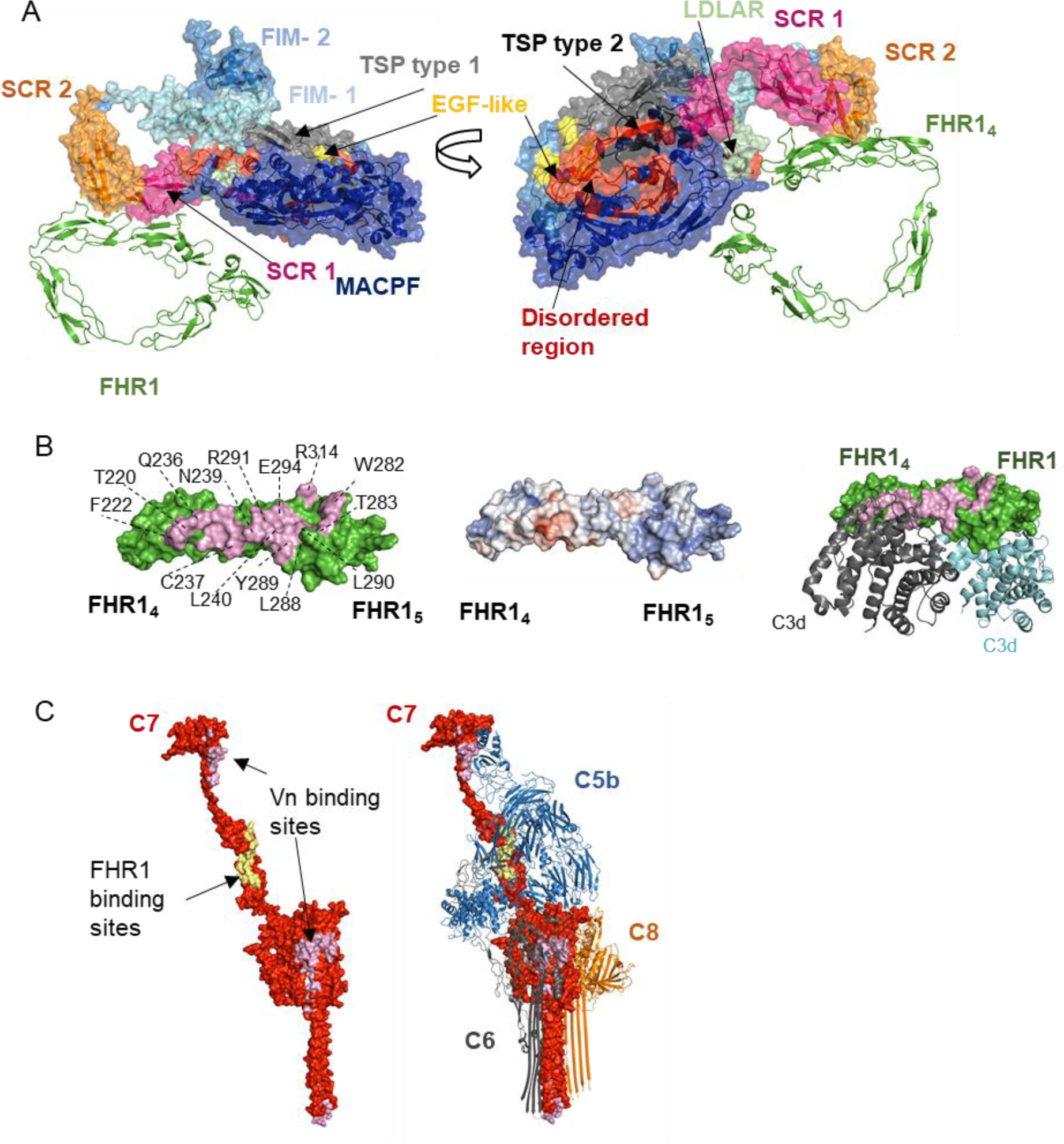
Interactions between FHR1 and C7 predicted by AlphaFold-Multimer. **A.** FHR1_4-5_ interacts with SCRs 1-2 domains of C7 according to top-ranked AFM model. The colours of nine C7 domains, shown as surface, match the colour of their legends and FHR1 is shown as cartoon in green. **B.** Binding interfaces in FHR1 of the model presented in A (on the left) with electrostatic potential surface (in the middle, electropositive residues in blue and electronegative in red). Some of the residues forming the binding region of FHR1 are labelled (interface shown in light pink). The binding region in FHR1_4-5_ partly overlaps with C3d binding region (on the right). (Experimental structures of C3d/FHR1_4-5_, PDB 3rj3). **C.** FHR1 and vitronectin binding to C7 would interfere with C5b-7 complex formation. The C7 interacting interfaces with FHR1 and Vn are marked in pale yellow and purple, respectively, on the experimental C7 structure in the soluble MAC (PDB 7nyc). The complex C5b-8 is shown with C7 as a surface and C5b, C6, C8 as cartoon.

According to superimposition of the AFM model with the model derived from cryoEM reconstructions of C7 in soluble MAC [61], the binding sites do not overlap with C8 binding sites in the C5b-7 complex (Figure 5 C). However, FHR1 binding sites overlap partially with C5b-C7 interfaces, which suggest that FHR1 binding to C7 would prevent the formation of the C5b-7 complex (Figure 5 C). It is important to highlight that FHR1-MAC interacting interfaces overlap partly with the C3d binding region.

This mechanism of action is similar to those observed for clusterin and vitronectin, the most important soluble regulators of the MAC. These proteins bind to TCC proteins and according to affinity assays on lipid bilayer, Vn prevents the binding of C5b-7 to the membrane, while clusterin binds to it without blocking the membrane association site [62]. These multifunctional proteins have large disordered regions and, due to their flexibility, the tertiary structure could not be experimentally determined. Therefore, the molecular mechanism by which Vn and clusterin recognize and block the MAC is not well understood [61]. Here, we analysed whether AFM can predict interactions between Vn and C7 and compared them with FHR1. The top ranked models across different runs with several recycle numbers show a clear interaction interface between C7 and Vn (Figure 5 C, Supplementary Figure 6 B). Different potential binding interfaces were found, which agrees with a previous study in which several Vn molecules were identified cross- linked in soluble MAC by combining cryoEM and cross-linking mass spectrometry (MS) [61].

According to our models, none of the interfaces were predicted in the heparin-binding region (362-395) of Vn, as was initially postulated [63] and contradicted later [64]. In the AFM model, the largest interface was located in a part of hemopexin 3-4 domains of Vn and a region of C7 MACPF (around residues 166-306) with 27 and 32 residues interacting, respectively, 17 strong hydrogen bonds, 4 hydrophobic interactions, 4 salt bridges and 2 electrostatic interactions. A hydrophobic cluster was identified with an area of 965.9 Å_2_ comprising 20 residues and 2.2 contacts/residue. Residues involved in the hydrophobic cluster are Leu 319, Leu 320 and Ile 316 in vitronectin and Ile 263 and Leu 184 in C7 (Supplementary Figure 6 C). The key residues in the interface are Lys 420, Ile 263 and Ser 185 in C7, and Arg 330 and Leu 319 in Vn, involved in salt-bridges and hydrophobic interactions.

The second interface between Vn and C7 is located in parts of hemopexin 1-2 of Vn and the first factor I-like membrane attack complex domain (FIM 1) of C7 with 17 and 18 interacting residues, respectively, forming 14 strong hydrogen bonds, 2 hydrophobic interactions, 5 salt-bridges and 3 additional electrostatic interactions (Supplementary Figure 6 B). Some key residues in the interface are Arg 712, Arg 753, Met 717 in C7, and Asn 169, Trp 322, Asp 232 in Vn, involved in hydrogen bonds, salt-bridges and hydrophobic interactions.

Recently, it was suggested that FIM 1 of C7 is responsible for the adaptation of C5b to enable the complete MAC assembly, since it can reorient the C5b C345C domain and triggers the accommodation of macroglobulin domains MG4 and MG5 of C5b to recruit C8β [61]. According to our models, interaction of Vn with FIM1 of C7 would avoid the binding of C7 to the C5b C345C domain in C5b6, hindering its activation. Likewise, the second Vn-binding site in C7, MACPF, overlaps in a small part with the region that undergoes a conformational change to form transmembrane β-hairpins and insert into the membrane. Binding of Vn to this site might avoid the conformational change of C7 to penetrate the membrane and interfere with C6 binding, preventing C5b-7 complex formation and membrane insertion (Figure 5 C, Supplementary Figure 6 D). New binding sites between Vn and the C5b-7 complex might be formed after dramatic conformational rearrangements of C7, C5 and C6 that are not predicted by AF2. Moreover, there are additional binding sites in C5b and C6 in the complex C5b-7 with Vn shown by combining cryoEM and cross-linking MS [61] which might be important for soluble MAC clearance and regulation of C5b-8 complex formation.

Although the binding sites on C7 for FHR1 and the hemopexin 1-2 and 3-4 domains of vitronectin are different, the mechanism of action would be similar, avoiding the formation of a C5b-7 complex. Reports about correlation between protein-protein interactions derived from the structure and their binding affinity are limited, however, a moderate correlation was shown for features such as number of H-bonds and geometric complementarity [65]. The predicted interacting interfaces with C7 are larger for Vn than for FHR1 and could explain why vitronectin is one of the main regulators found associated to soluble MAC and might correlate with a higher binding affinity.

#### 2.3.3 Interactions between FHR1 and C9

The binding of C9 to C5b-8 is the kinetic bottleneck of MAC formation, followed by rapid unidirectional polymerization of additional C9 to build a circular pore [59].

The models for the analysis of the interaction between C9 and FHR1 were predicted by AFM with several recycle numbers, amber relaxation and with or without templates. Top ranked models suggest interactions between C9 and FHR1 with interfaces located in domains FHR1_1-2_, FHR1_5_ and a small region in FHR1_4_, involving 10, 14 and 3 residues, respectively (Figure 6 A, Supplementary Figure 7 A). The FHR1_1-2_ binding interface forms 10 hydrogen bonds and 8 hydrophobic interactions, while the FHR1_4-5_ binding interface forms 17 hydrogen bonds, 9 electrostatic interactions and 4 hydrophobic interactions. The key residues in the top-ranked model predicted by AFM and identified by MM-GBSA implemented in the HawkDock server were Phe 66, Glu 294, Tyr 34, Arg 302, Trp 282, Glu 297, Tyr 57, Gln 242, and Ser 65. Both binding sites are located in a region of the C9 MACPF domain (Figure 6 A, B). As mentioned above, it is still difficult to predict the correct orientation of FHR1 domains, therefore, FHR1 might interact simultaneously or independently through several binding regions with C9.

**Figure 6.**
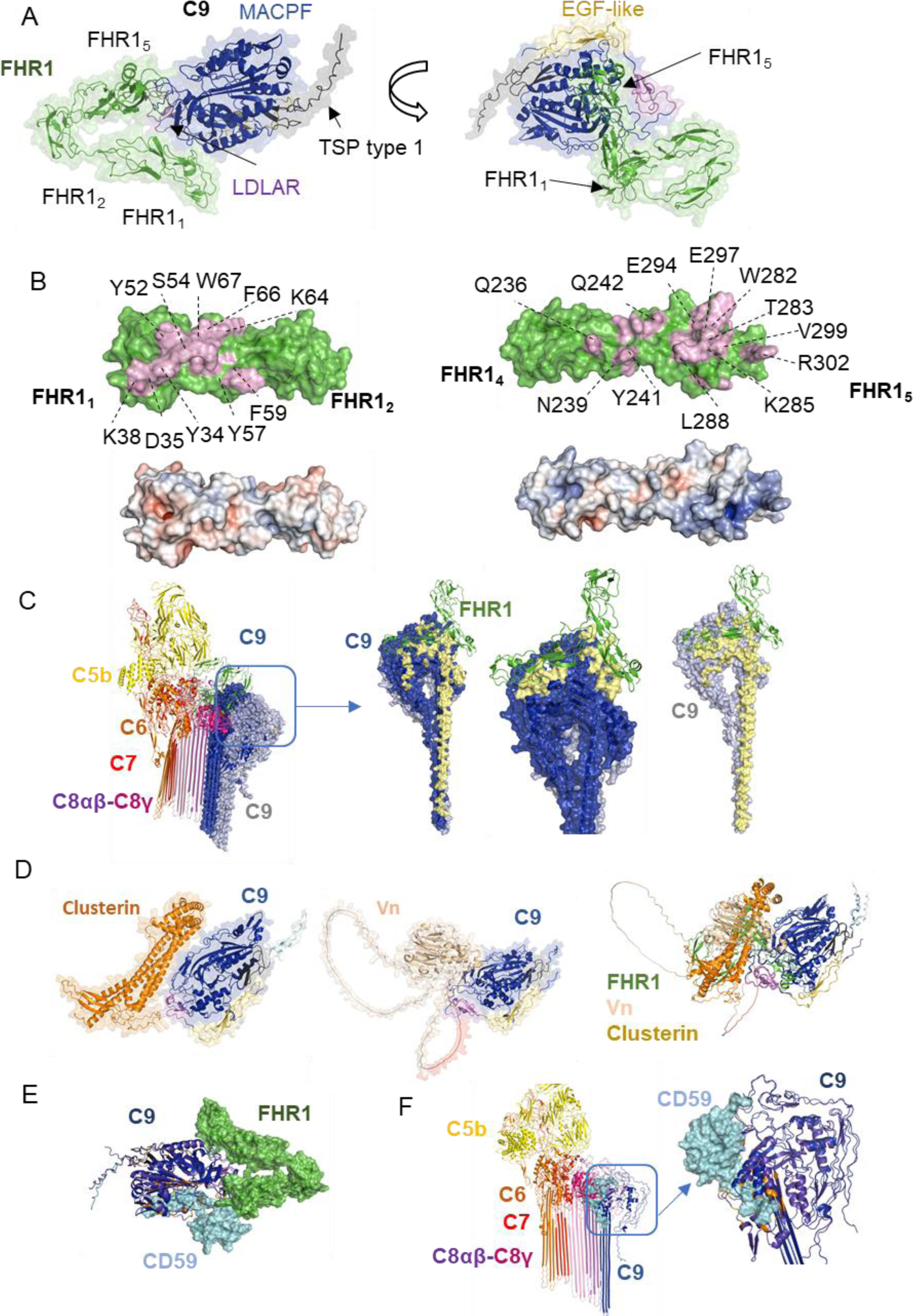
Interactions between FHR1 and C9 predicted by AlphaFold Multimer **A.** FHR1_1-2_ and FHR1_5_ interact with C9 MACPF domain according to the AFM model. FHR1 is shown in green and the colours of C9 domains match the colour of their legends. **B.** Binding interfaces of the model presented in A with electrostatic potential surface (electropositive and electronegative residues in blue and red, respectively). Some of the residues forming the binding region are labelled in FHR1. **C.** Interactions of vitronectin and clusterin with C9 predicted by AFM and superimposition of all models including the FHR1 complex, suggest similar binding sites (only one of the top-ranked models is presented). **D.** FHR1 binding to C9 avoids C9 polymerization. Superimposition of C9 and FHR1 model with experimental structure of MAC (PDB 7nyc) is shown with a zoom in of interfaces between C8α -C9 and C9-C9, contrasted with binding sites of FHR1 (green) to C9 (blue and grey) in pale yellow. **E.** Interactions between membrane-anchored MAC regulator CD59 (cyan) and C9 (dark blue) predicted by AFM are located in a buried region of soluble C9. The binding site is different from the predicted binding site for FHR1 (green). **F.** Superimposition of the AF2 model and experimental structure of MAC (PDB 7nyc) shows binding sites of CD59 in a hydrophobic pocket where unfurling of transmembrane β-hairpins occurs (binding sites in orange, soluble C9 and MAC C9 in purple and blue, respectively).

The binding sites in FHR1_1-2_ coincide with the dimerization interface of FHR1. Therefore, similar to C7, binding to C9 would compete with the dimerization of FHR1. The C9-binding interface in FHR1_4-5_ differs from the binding interface to TED domain of C3 (or C3 fragments). However, FHR1 cannot bind both molecules simultaneously due to steric hindrances and clashes (Figure 6 B). To explore the effect of FHR1 and C9 interaction on MAC formation, our models were superimposed with a structure of the C5b-8 and C5b-9 complex. The best model predicted by AFM indicates FHR1 binding to a C9 MACPF region which does not undergo conformational changes upon C9 binding to C5b-8, indicating that FHR1 can also bind to monomeric C9 in MAC. Furthermore, FHR1 would block C9 polymerization, since the binding region with the C9 monomer coincides with the C9-C9 polymerization interface (Figure 6 C). The interaction between FHR1 and native/soluble C9 would block binding to the C5b-8 complex (Figure 6 C).

Clusterin and vitronectin can bind to C9, and block its polymerization and pore formation [64, 66]. Clusterin binds to precomplexes C5b-7, C5b-8, C5b-9 but not to C5b6, and it binds to individual TCC proteins C7, C8β and C9, but not C5, C6, C8α and C8γ [67]. Here, we also included a model of these complement regulators in complex with C9 and compared them with FHR1. According to the models predicted by AFM, Vn interacts with C9 mainly by a hydrophobic region which involves Trp 424, Trp 322 and Tyr 420 in Vn and Ile 203, Trp 197 and Arg 116 in C9, along with electrostatic interactions and hydrogen bonds (Supplementary Figure 7 B). On the other hand, clusterin interacts with C9 mainly by electrostatic interactions (including salt bridges) which involve Asp 178, Asp 174 and Glu 189 in clusterin and Arg 145, Arg 183 and Lys 114, along with hydrogen bonds (Supplementary Figure 7 C, D). Some of the key residues in the binding interface between C9 and Vn are some of those involved in hydrophobic interactions, while key residues in the interface with clusterin correspond to those involved in electrostatic interactions (Figure 6 D). These models are congruent with previous experimental results indicating that Vn binds to a hydrophobic region of C9, while clusterin interacts with a charged patch in C9 [61].

The binding interfaces of clusterin, vitronectin and FHR1 are located in a similar region of the C9 MACPF domain where they interfere directly with binding to C5b-8 complex (Figure 6 D).

However, some of the top ranked models of clusterin showed additional binding sites (Supplementary Figure 7 E, F, G), which indicates that there are different interfaces on C9 for these regulators as suggested previously [61]. Furthermore, monomeric or polymerized C9 can compete with clusterin for the binding to C9 [68], which is congruent with our models of C9/FHR1 and C9/clusterin complexes, indicating that the binding interface coincide with the polymerization C9-C9 interface.

The regulator CD59 binds C8α and C9 in the C5b-8 or C5b-9 complexes during MAC assembly, preventing C9 binding and C9 polymerization, respectively [58, 69]. Although the experimental structures of CD59 complexes with the complement components have not been reported, binding sites have been identified by peptide screening [69]. Therefore, we performed a prediction using AFM to evaluate whether it is congruent with the experimental data and compared it with FHR1 binding sites. A hydrophobic pocket in C9 was identified previously as the primary binding site for CD59 by docking models and mutagenesis studies [69–71], which is congruent with our AFM prediction (Figure 6 E, Supplementary Figure 7 H). This region is buried within C9 β-hairpins and undergoes a conformational change when C9 binds to the membrane-bound C5b-8 or C5b-9 complexes. Since CD59 cannot bind soluble C9, it should bind during the conformational rearrangement of transmembrane helices, and it might also prevent a complete conformational change (Figure 6 F). Furthermore, the CD59 binding site could generate clashes with the C8γ subunit, therefore CD59 would not bind to the first C9 molecule recruited to the C5b-8 complex. Since the recruitment and insertion of the first C9 was recognized as the rate-limiting step in MAC formation [59], CD59 can act initially by binding to C8α chain to prevent bilayer perforation and C5b-9 complex formation. Once this complex is formed, CD59 binds to subsequent C9 in the complex to prevent further oligomerization. According to our AFM models, CD59 and C9 interact mainly by hydrophobic interactions with a binding region located in the transmembrane helices of C9, which include the peptide VSLAFS identified in C9 by experimental approaches [69]. Key residues in the interface include Phe 391, Phe 400, Phe 455, Phe 450, Val 452 in C9 and Phe 119, Leu 115, Leu 120, Trp 124, Pro 118 in CD59. CD59 might interfere with FHR1_5_ interaction with C9 by steric hindrances but its binding site does not overlap with the binding region of FHR1_1_. Therefore, our models indicate that CD59 might not compete with FHR1 (Figure 6 E).

Although the confidence scores of the models between complement regulators such as clusterin, vitronectin, CD59 with C7 or C9 (inter-chain predicted alignment error) are lower than those for FHR1/C3d complexes, binding interfaces predicted by AFM are in agreement with experimental observations. Lower confidence scores were also obtained for all models between FHR1 in complex with C5, C7 or C9 than those for FHR1/C3d (Supplementary Table 1, Supplementary Figures 5, 6 A, 7 A), although we have experimentally measured a higher K_d_ for C5, C9 and C7 than for C3d in the case of MFHR13, closely related to FHR1 [13]. Our models suggest that FHR1 competes with itself due to the dimerization interface for binding to C9. Moreover, it would compete with other ligands such as C3, C3b, C3d for binding to C5, C7 or C9. These findings might explain why FHR1 needed molar excess in respect to TCC proteins to inhibit MAC on sheep erythrocytes [13].

As Alphafold-Multimer version 1 (AFM-v1) results frequently in clashes between protein chains, ColabFold also implemented an improved version, AFM version 2 (AFM-v2) [72]. All results presented here were obtained using AFM-v2. However, we compared both versions to predict FHR1 in complex with C5, C7, C9 and C3d using different recycle numbers. Increasing the recycle number has a severe impact in AFM-v1 confidence scores compared with AFM-v2.

AFM-v1 and AFM-v2 correctly predicted binding interfaces in the complex FHR1/C3d, but we always observed higher confidence model scores for AFM-v1 prediction (pLDDT, pTM and iptm), including FHR1 in complex with C5, C7 and C9 (Supplementary Table 1). However, prediction with AFM-v1 always led to clashes in the case of the C5/FHR1 complex, and in some cases also for C7 and C9. We included an additional top-ranked model obtained with AFM-v1 that was ranked as the best predicted for the C9/FHR1 complex in which we did not detect clashes and suggest potential binding interfaces in a different region of C9 compared with AFM- v2 (Supplementary Figure 8 A). In that case, FHR1_1_ binds to a region of C9 MACPF that undergoes a conformational change to insert into the membrane, indicating that FHR1 would interfere with the conformational change of C9 upon C5b-8 binding (Supplementary Figure 8 B). The models from the two different versions are not contradictory, as several potential binding sites of FHR1 are possible for these TCC proteins, as in the case of clusterin and vitronectin.

Our models predicted binding interfaces in FHR_1-2_ or FHR1_4-5_ with C5, C7 and C9. It is important to mention that MFHR13 includes FHR1_1-2_ and FH_19-20_, which differ from FHR1_4-5_ only by two residues (SV in FH_20_ and LA in FHR1_5_). Nevertheless, we have not detected binding of FH to these proteins in wet lab experiments [13]. These differences have important implications for the action mechanism of both proteins; the SV motif (FH) binds efficiently to sialic acids contrary to the LA motif (FHR1) [10]. These differences might explain previous results where MFHR13 protected C3b-opsonized sheep erythrocytes from convertase-independent C5 activation and MAC formation more efficiently than FHR1 [13], as MFHR13 (FHR1_1-2_:FH_1-4_:FH_13_:FH_19-20_) would bind more efficiently to the sialic acid-rich surface of sheep erythrocytes, regulating MAC formation on the cell surface. Furthermore, FH binds more efficiently to heparin-C3b while FHR1 binds better to a heparin-C3d combination [73]. Although in none of our models Leu 290 or Ala 296 were identified as key residues in the binding interfaces of FHR1, the effect of these motifs (SV and LA, on FH and FHR1, respectively) on the binding capacity to C3b/C3d, C5, C7 and C9 should be experimentally investigated, in order to validate the binding interfaces in FHR1_4-5_ and to elucidate if the inability of FH to bind to C5, C7 and C9 depends on the sequence of FH_20_ or on the conformation of FH in fluid phase.

The role of FHR1 as a MAC complement regulator has been controversial. However, it has been proposed that FHR1 acts either as complement regulator or “FH antagonist” depending on the context [74, 75]. For instance, the knockout of the murine FHR1 homolog resulted in mice with severe sepsis, acute kidney injury and alternative pathway overactivation in response to LPS challenge [76]. Therefore, under physiological conditions, FHR1 could promote complement opsonization on damaged host cells to facilitate phagocytosis, while preventing MAC formation and restricting inflammation. In case of complement overactivation during disease FHR1 might act as a MAC regulator.

## 3. CONCLUSION

We used AF2 and AFM to predict the structures of synthetic and native human complement regulators and provide insights into their molecular interactions with complement components. Differences in MFHR1 and MFHR13 structures explain differences in alternative pathway regulation observed before experimentally, supporting these results [13]. However, these models should be interpreted with caution, since domain orientation is still difficult to predict. AFM successfully predicted binding interfaces of FHR1 and the synthetic regulators to C3 ligands and predicted binding sites of FHR1 with TCC proteins (C5, C7 and C9) located in FHR1_1-2_ and FHR_4-5_, which indicates that FHR1 could block C5 activation, C5b-7 formation and C9 polymerization, with a similar action mechanism to Vn, clusterin.

We generated various hypotheses based on AF2/AFM predictions in order to understand the molecular mechanism of complement regulation, with emphasis on MAC inhibition. These may serve as a basis to design straightforward experiments using a rational approach to identify binding interfaces of MAC proteins C5, C7, and C9 with natural regulators (FHR1, clusterin or Vn) or synthetic regulators (MFHR1 or MFHR13). In this context, AF2 and AFM can be powerful tools to unravel complement regulatory mechanisms and design novel complement therapeutics. For example, several pathogens have evolved to evade an attack of the complement system, among them *Neissseria meningiditis, Borrelia miyamotoi*, *Yersinia* and *Salmonella* species which can recruit FH or Vn to their surface [77, 78]. Prediction of such interactions via AF2/AFM would simplify unveiling mechanisms of immune evasion and might lead to targeted treatments.

## 4. METHODS

### 4.1 Prediction of protein structures and protein-protein interactions

The sequences of complement proteins were retrieved from UniProt. To predict protein structures, we used Colabfold [29], which is based on the AlphaFold2 (AF2) algorithm to predict monomers [27] and AlphaFold-Multimer (AFM) [28] to predict protein complexes. ColabFold uses MMseqs2 (Many-against-Many sequence searching) instead of AF2 homology search (HMMer and HHblits), which significantly accelerates the prediction, while matching the prediction quality [29].

We used local ColabFold, a free open-source software, which allows ColabFold to run on our local machines instead of Google Colaboratory. We ran ColabFold in Linux with Nvidia GPU V100 (CUDA 11.1 was installed). All models were run using amber relaxation, which is important for the positioning of the side chains and to avoid steric clashes. The number of recycles was modified from the default settings. This is the number of times the prediction is fed through the model which can significantly improve the quality [29]. Although 3 is used by default, we also included 6, 8, 12 and 20 recycles. We used templates for the protein complexes in PDB70 and compared them with models generated without templates.

ColabFold implemented two algorithms to predict protein-complexes based on AlphaFold-multimer (AFM) version 1 and 2 [28, 29]. AF2/AFM computes five models for every run and we analysed the models with the best score across all runs with variable recycle numbers. AF2 model quality is evaluated using two metrics, the predicted Local Distance Difference Test (pLDDT) and predicted Aligned Error (PAE). For protein complexes, inter-chain PAE and interface pTM score (iptm) was used to evaluate the accuracy of interfaces and model confidence. Inter-chain PAE plots are shown in supplementary data. High and low prediction qualities are indicated in blue (values close to 0) and red (values close to 30), respectively.

The quality of the synthetic regulator (MFHR1 and MFHR13) models were additionally evaluated using Ramachandran Plot in the PROCHECK module implemented in the SAVES server to analyse stereochemical quality (https://saves.mbi.ucla.edu/), and QMEANDisCo [35] (https://swissmodel.expasy.org/qmean/).

### 4.2 Other tools

We used Pymol 2.5.4 (Schrödinger, LLC, New York, NY, USA) to visualize and analyse the 3D protein structures including analysis of interactions between proteins (using the plugins show_contacts, list_contacts), calculation of electrostatic surface potential, and root-mean-square deviation of atomic positions (RMSD) to compare the predictions with experimental structures by structural alignment (alignment /superposition plugin). The experimental protein structures were retrieved from the RCSB protein data Bank (https://www.rcsb.org/). The following PDB structures were used for analysis: PDB 2wii (complex FH1-4/C3b), PDB complex mini-FH: C3b (5035), PDB 2a73 (C3), PDB 2xqw (C3d/FH19-20), PDB 4muc, 3rj3 (C3d/FHR14-5), PDB 7nyc (soluble MAC with 3C9), PDB 2kms (FH12-13), PDB 3sw0 (FH18-20), PDB 2win (C3 convertase C3bBb) PDB 4ont (C3d/sialic acid/FH19_20), PDB 3zd2 (dimer FHR11-2) PDB 4e0s (C5b6).

Intermolecular interactions, bumps and clashes were also analysed using the BIOVIA Discovery Studio Visualizer 21.0.0 (Biovia Dassault Systèmes, San Diego, USA). Other model features such as hydrophobic clusters were analysed using the ProteinTools web server [79].

The intrinsic disorder regions were identified with the following web tools: IUPred2A [80], DISOPRED3 [81], fMoRFpred [82], DFLpred [83] and TransDFL [84].

Binding free energy of the best models was calculated using the molecular mechanics/generalized Born surface area (MM/GBSA) method implemented in the HawkDock server to identify key residues in the binding interfaces [85].

## ACKNOWLEDGEMENTS

This work was supported by the Mathias-Tantau-Stiftung (to NRM) and the Deutsche Forschungsgemeinschaft (DFG, German Research Foundation) under Germany’s Excellence Strategy EXC-2189 (CIBSS; to RR). We thank the German Network for Bioinformatics Infrastructure (de.NBI) for the computational resources, Andres Posada Moreno for his help with setting up the servers, Anne Katrin Prowse for proof-reading of the manuscript, and Michal Rössler for her help with the graphical abstract.

## Declaration of competing interests

All authors declare to have no competing interests.

## Authorship contribution statement

NRM: conceptualization, methodology, formal analysis, writing-original draft preparation. JP: conceptualization, writing- original draft preparation. ELD: funding acquisition, writing- reviewing and editing. RR: conceptualization, funding acquisition, writing- reviewing and editing. All authors read and approved the final manuscript.

## Supplementary information

**Supplementary Figure 1.**
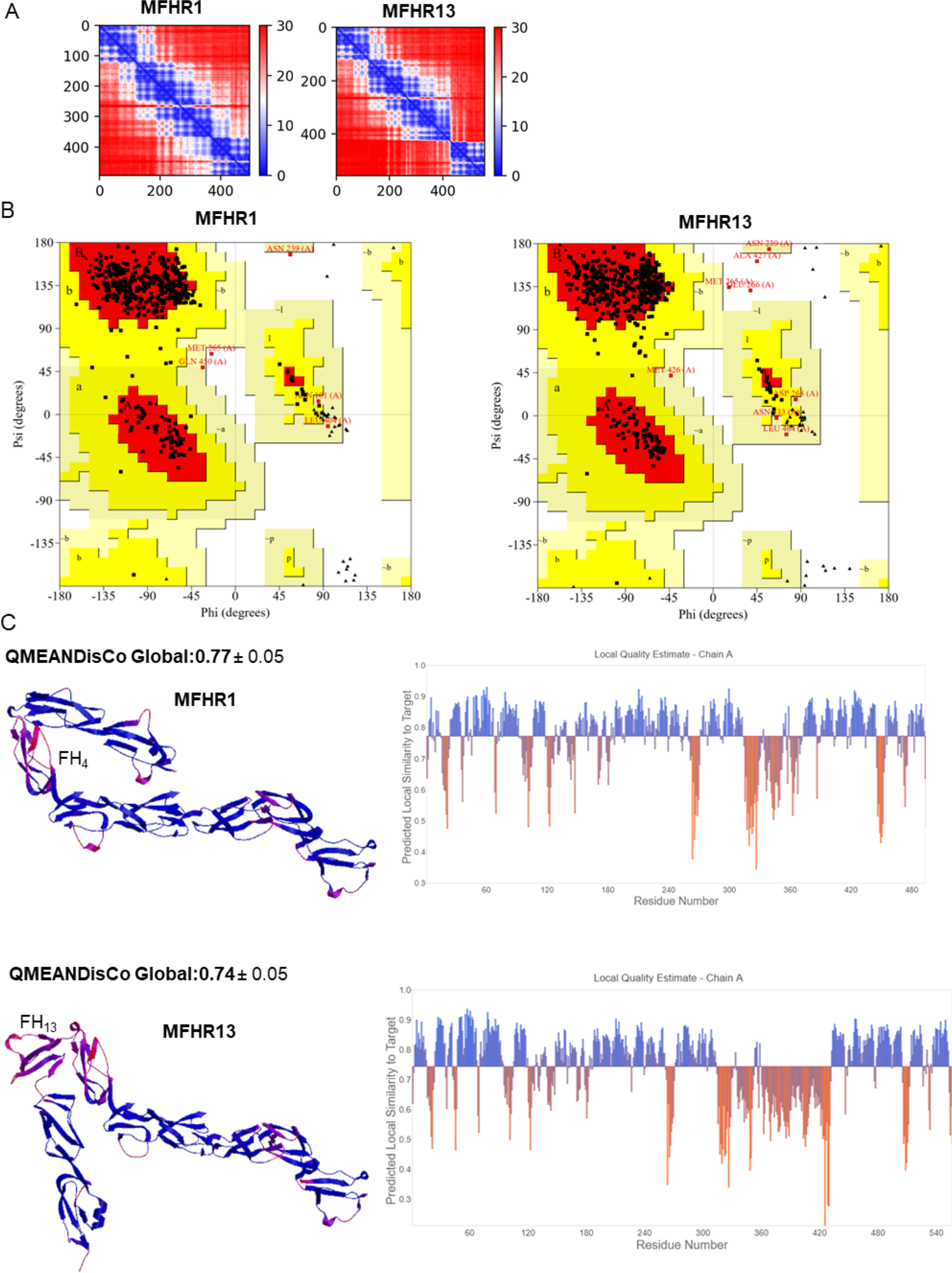
Quality assessment of MFHR1 and MFHR13 structure models. **A.** Predicted Aligned Error (PAE) plot for the best-ranked model structures of MFHR1 and MFHR13 by AlphaFold2 shown in figures 1A and 1B, respectively. The orientation between domains is not predicted with a high confidence. **B.** Ramachandran plots calculated by PROCHECK. On the left: MFHR1 shows 87.8% residues in most favoured regions [A, B, L, in red], 11% residues in additionally allowed regions [a,b,l,p], 0.7% residues in generously allowed regions [̴ a, ̴ b, ̴ l, ̴ p] and 0.5% residues in disallowed regions. On the right: for MFHR13 there are 89.3% residues in most favoured regions [A, B, L, in red], 9% residues in additional allowed regions [a,b,l,p], 1.1% residues in generously allowed regions [̴ a, ̴ b, ̴ l, ̴ p] and 0.6% residues in disallowed regions. Outliers in disallowed regions are located in the 13-19 linker. **C.** Model quality assessed by QMEANDisCo. The global scores were 0.77 and 0.74 for MFHR1 and MFHR13, respectively. Local quality for each amino acid residue visualized in the structure model is shown on the left, while local quality scores of each amino acid residue is shown on the right. Low and high confidence regions are shown in a scale from red to blue.

**Supplementary Figure 2.**
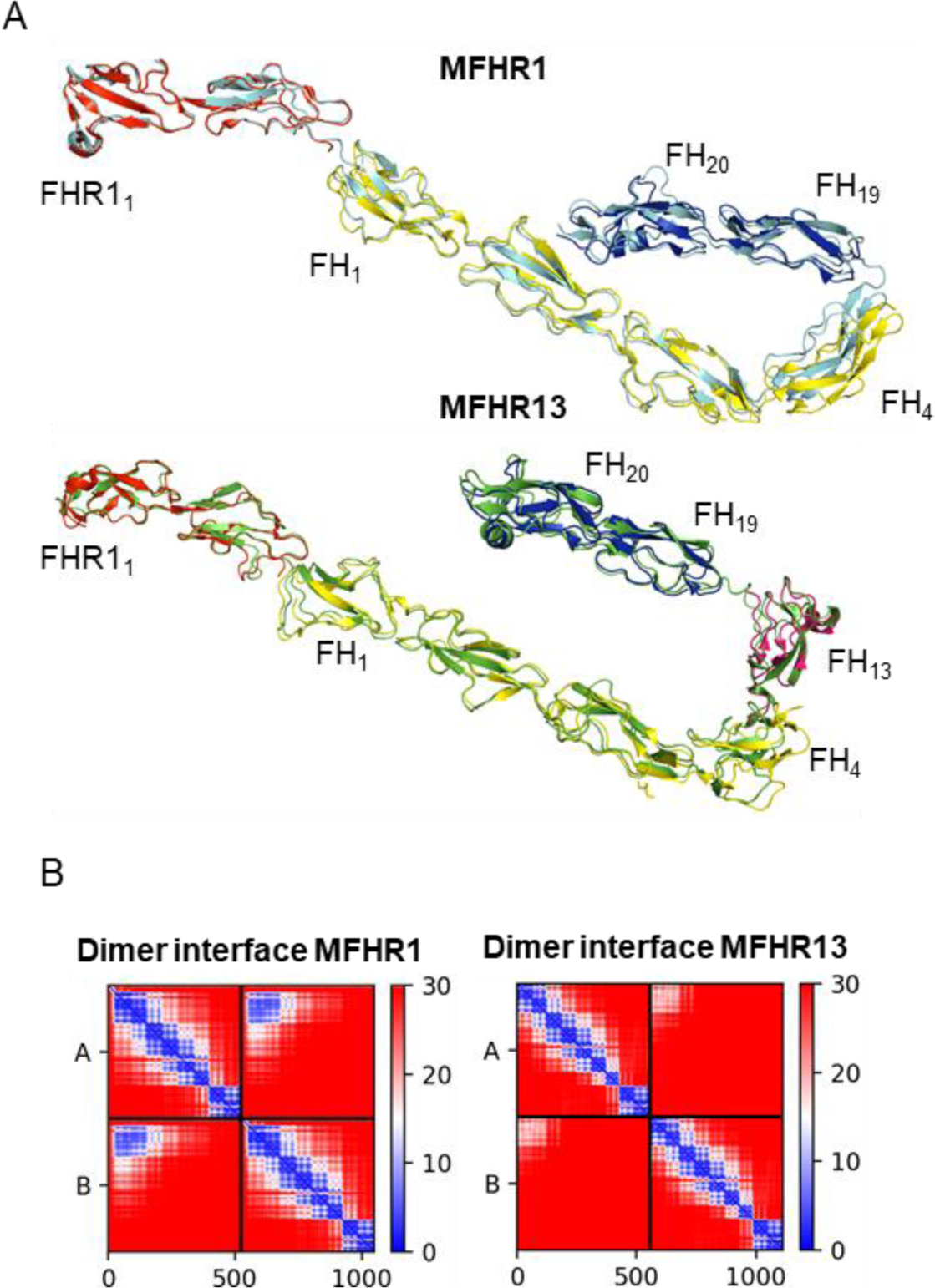
Additional quality assessment of MFHR1 and MFHR13 structure models and dimerization interfaces predicted by AlphaFold 2 and AlphaFold-Multimer (AFM), respectively. **A**. Structure models of MFHR1 (light blue) and MFHR13 (green) were superimposed with experimental structures. FHR1_1-2_ (PDB 3zd2) in red with a root-mean-square deviation (RMSD) of 0.595 Å and 0.627 Å for MFHR1 and MFHR13, respectively. FH_1-4_ (PDB 2wii) in yellow with RMSD = 1.266 Å and 1.376 Å for MFHR1 and MFHR13 respectively. FH_19-20_ (PDB 2xqw) in dark blue with RMSD= 1.260 Å and 1.209 Å for MFHR1 and MFHR13, respectively. FH13 (PDB 2KMS) in pink with RMDS = 0.939 Å for MFHR13. **B.** Inter-chain Predicted Aligned Error (PAE) plot for the best-ranked model structure of MFHR1 and MFHR13 dimerization by AFM shown in figure 1 C.

**Supplementary Figure 3.**
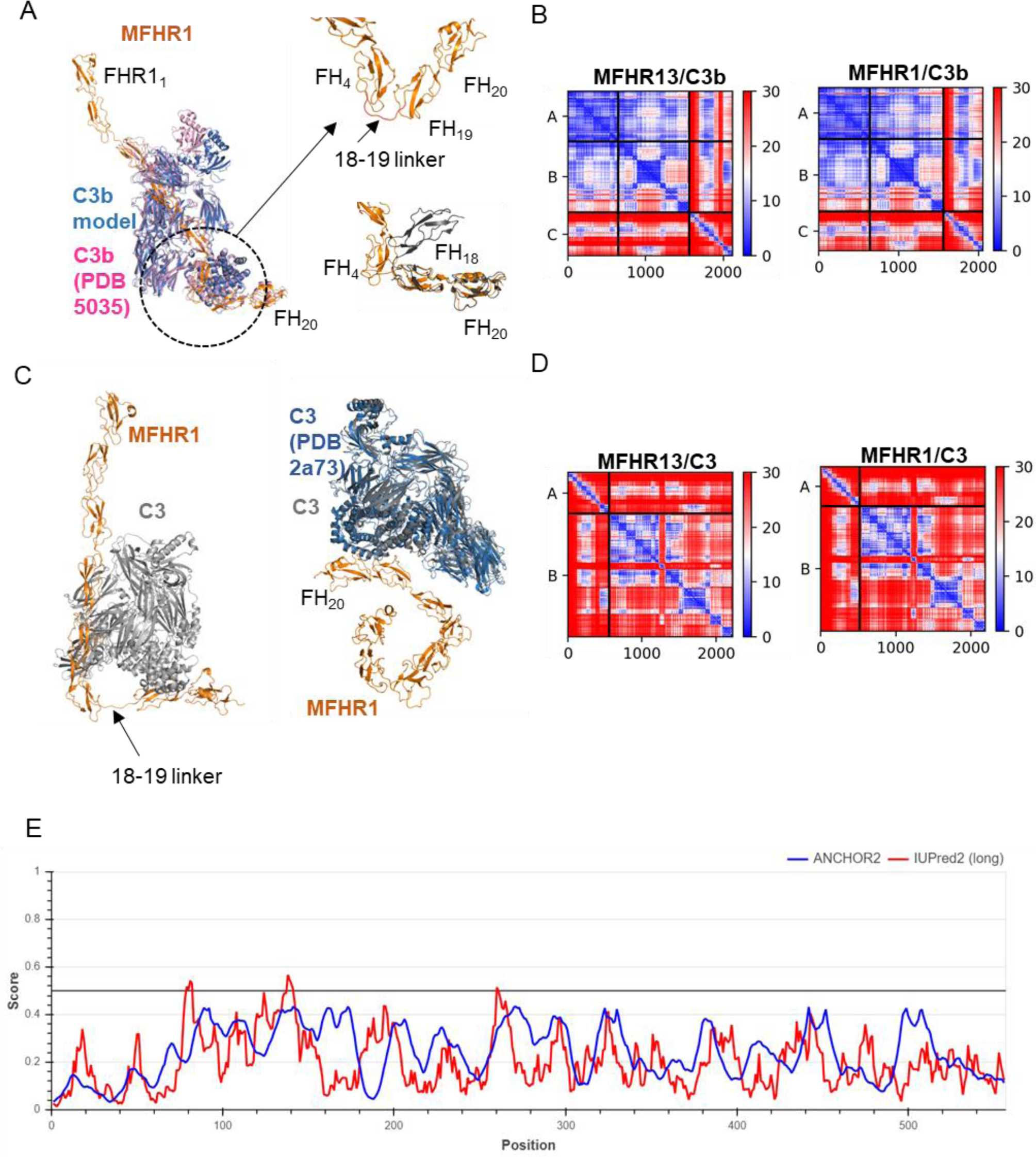
Quality assessment of complex models between FHR1, MFHR1 or MFHR13 in complex with C3 or C3b. **A.** Structure of MFHR1 (orange) in complex with C3b (blue) predicted by AFM. Superimposition of the model with the experimental structure of mini-FH/C3b (PDB 5035, pink) with RMSD = 1.605 Å. On the right the orientation of FH_4_, FH_19_ and FH_20_ domains in MFHR1 in complex with C3b and the experimental structure FH_18-20_, which shares the 18-19 linker (PDB 3sw0, black). **B.** Inter-chain Predicted Aligned Error (PAE) plot for the best-ranked models of MFHR13/C3b and MFHR1/C3b complexes predicted by AFM. The scores for MFHR1/C3b (12 recycles) were pLDDT = 86.5, ptmscore = 0.823 and iptm = 0.795 (shown in Figure 2 A) and for MFHR13/C3b (6 recycles) the scores were pLDDT = 84.7, ptmscore = 0.79 and iptm = 0.754. **C.** Structure of MFHR1 (orange) in complex with C3 (grey) predicted by AFM. The extended conformation of the 4-19 linker (18-19 linker in FH) makes the interaction through both interfaces with one C3 molecule unreliable. On the right an alternative model with only one interacting interface in FH_19,_ and superimposition of C3 in the model with experimental structure PDB 2a73 (blue) with RMSD = 2.016 Å. **D.** Inter-chain Predicted Aligned Error (PAE) plot for the best-ranked models of MFHR13/C3 and MFHR1/C3 and complexes predicted by AFM. The scores for MFHR1/C3 (12 recycles) were pLDDT = 75.8, ptmscore = 0.592 and iptm = 0.497 and for MFHR13/C3 shown in Figure 2 B (12 recycles), pLDDT = 77.5, ptmscore = 0.561 and iptm = 0.452. **E.** Prediction of disorder regions in MFHR13 by IUPred2A (red), and the previous version of IUPred2A, Anchor (blue). A high score is associated with residues within a disordered region.

**Supplementary Figure 4.**
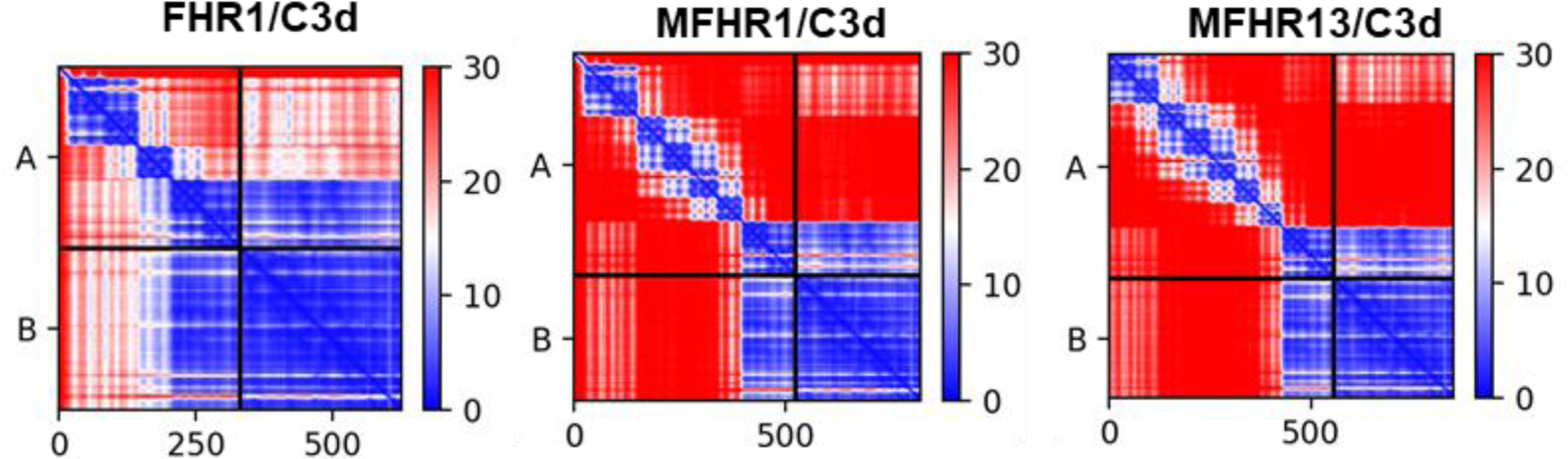
Inter-chain Predicted Aligned Error (PAE) plot for the best-ranked model of FHR1, MFHR1 and MFHR13 in complex with C3d predicted by AFM (shown in Figure 3). AFM scores for FHR1/C3d were: (12 recycles) pLDDT = 86, ptmscore = 0.751 and iptm = 0.87. For MFHR1/C3d: (12 recycles), pLDDT = 81.6, ptmscore = 0.555 and iptm = 0.849. MFHR13/C3d: (12 recycles) pLDDT = 84.5, ptmscore = 0.537 and iptm = 0.843.

**Supplementary Figure 5.**
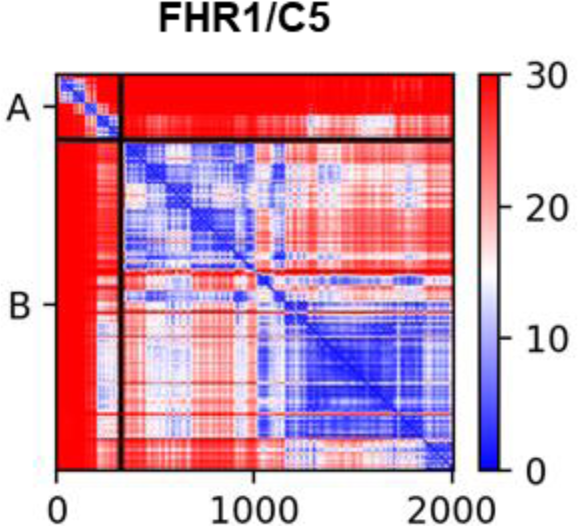
Inter-chain Predicted Aligned Error (PAE) plot for the best-ranked model of FHR1 in complex with C5 predicted by AFM, shown in Figure 4. AFM scores were pLDDT = 81.6, ptmscore = 0.693 and iptm = 0.384 (20 recycles).

**Supplementary Figure 6.**
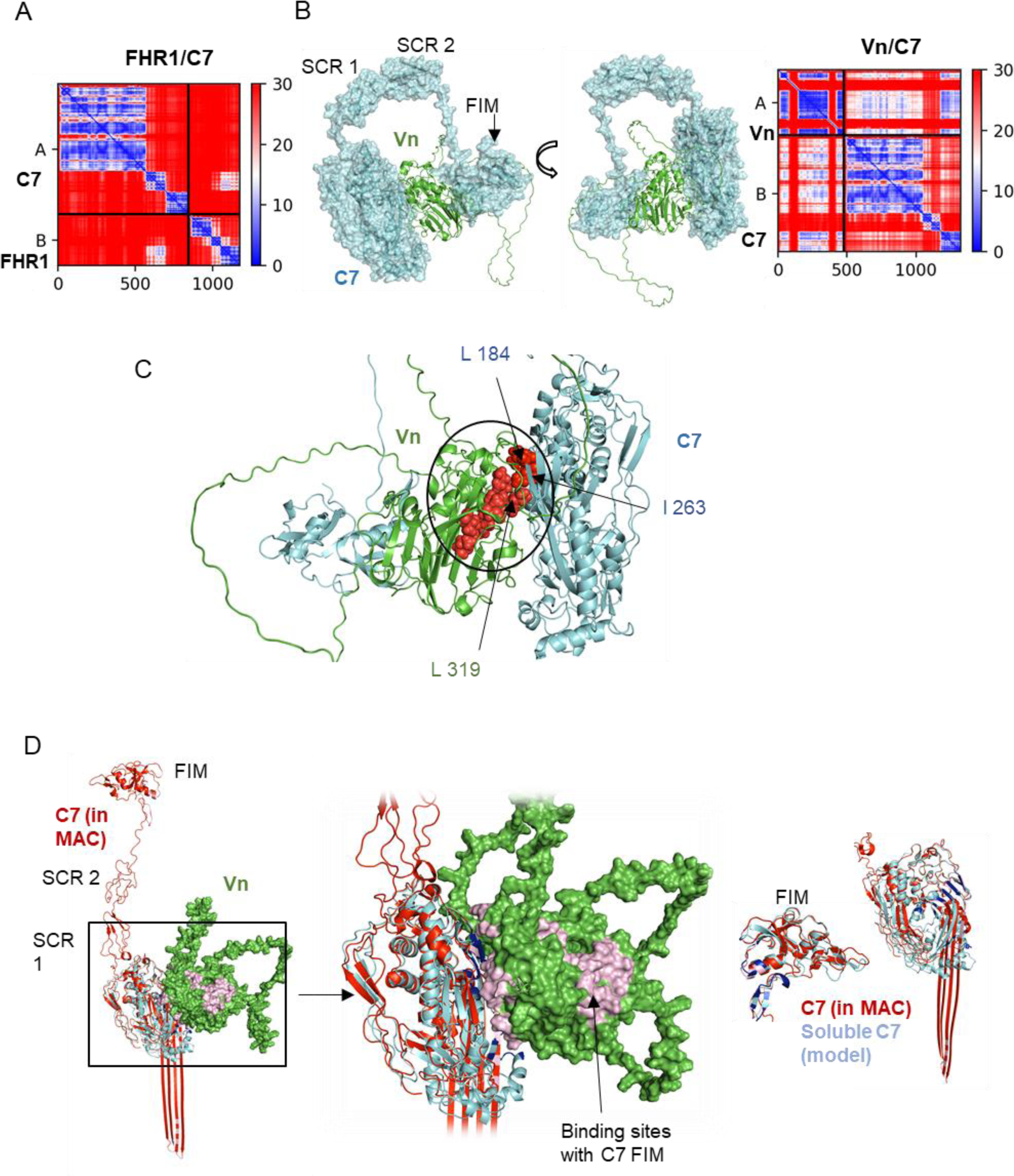
Interactions of C7 with FHR1 and vitronectin (Vn) predicted by AFM. **A.** Inter-chain Predicted Aligned Error (PAE) plot for the best-ranked model of FHR1 in complex with C7 with 20 recycles and pLDDT = 72.2, ptmscore = 0.512 and iptm = 0.303 (shown in Figure 5). **B.** Best-ranked model of vitronectin (green) in complex with C7 (blue) shows two binding interfaces located in a MACPF region and FIM domains in C7 (Figure 5 C). On the right the Inter-chain Predicted Aligned Error (PAE) plot with the following scores: pLDDT 71.9, ptmscore 0.568 and iptm 0.415. **C.** Hydrophobic clusters (red spheres) formed in the binding interface between the C7 MACPF region and Vn (key residues in the interface are labelled). **D.** Vitronectin binding to C7 would interfere with C5b-7 complex formation. Soluble C7 model (cyan) is superimposed with the experimental C7 structure (red) in the sMAC (PDB 7nyc). An additional interacting interface in Vn with C7 FIM domains is shown in pink. Conformational rearrangements of MACPF domain in C7 MAC superimposed with soluble C7 structure model are shown on the right.

**Supplementary Figure 7.**
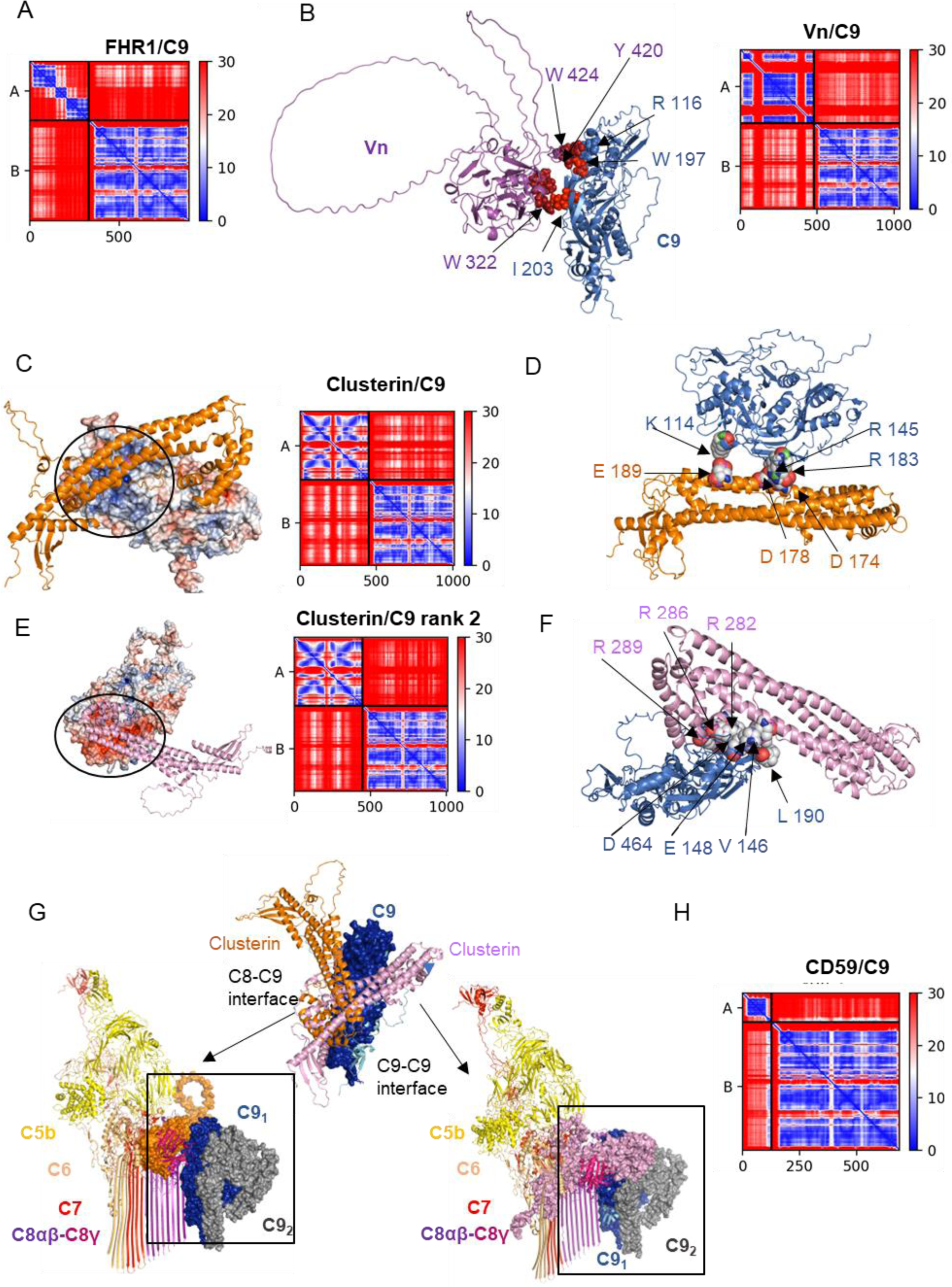
Interactions of C9 with FHR1, vitronectin, clusterin and CD59 predicted by AFM. **A.** Inter-chain Predicted Aligned Error (PAE) plot for the best-ranked model of FHR1 in complex with C9 (3 recycles) with the following scores: pLDDT = 74, ptmscore = 0.597 and iptm = 0.272 shown in Figure 6 A. **B.** Best-ranked model of vitronectin (purple) in complex with C9 (blue) (partially shown in Figure 6 D). Key residues in the binding interface involved in hydrophobic interactions are represented as red spheres. On the right the Inter-chain Predicted Aligned Error (PAE) plot (12 recycles) with pLDDT = 69.8, ptmscore = 0.576 and iptm = 0.287. **C.** Best-ranked model of clusterin (orange) in complex with C9 (partially shown in Figure 6 D). Electrostatic potential surface of C9 is shown (electropositive residues in blue and electronegative residues in red). An electropositive patch in C9 is interacting with clusterin. On the right the Inter-chain Predicted Aligned Error (PAE) plot (20 recycles) with pLDDT = 70.3, ptmscore = 0.543 and iptm = 0.229. **D.** Key residues in the binding interface between C9 and clusterin (orange) from model shown in C are involved in electrostatic interactions (shown as dots). Key residues were identified by MM-GBSA implemented in the HawkDock server. **E.** Second-best ranked model of clusterin (purple) in complex with C9. Electrostatic potential surface of C9 is shown (electropositive residues in blue and electronegative residues in red). An electronegative patch in C9 is interacting with clusterin. On the right the Inter-chain Predicted Aligned Error (PAE) plot (20 recycles) with pLDDT = 70.7, ptmscore = 0.545 and iptm = 0.224. **F.** Key residues in the binding interface between C9 and clusterin (purple) from model shown in E (shown as dots). Key residues were identified by MM-GBSA implemented in the HawkDock server. **G.** Superimposition of soluble C9 of models shown in C and E with C9 MAC (PDB 7nyc, dark blue as surface) indicates two opposing binding interfaces which match C8α-C9 and C9-C9 interfaces in MAC (in the middle). The experimental structure of partially assembled MAC (PDB 7nyc) is shown superimposed with each clusterin/C9 model complex (left, model shown in C; right, model shown in E). **H.** Inter-chain Predicted Aligned Error (PAE) plot for the best-ranked model of CD59 in complex with C9 predicted by AFM (12 recycles) with pLDDT = 72, ptmscore = 0.693 and iptm = 0.263 shown in Figure 6 E.

**Supplementary Figure 8.**
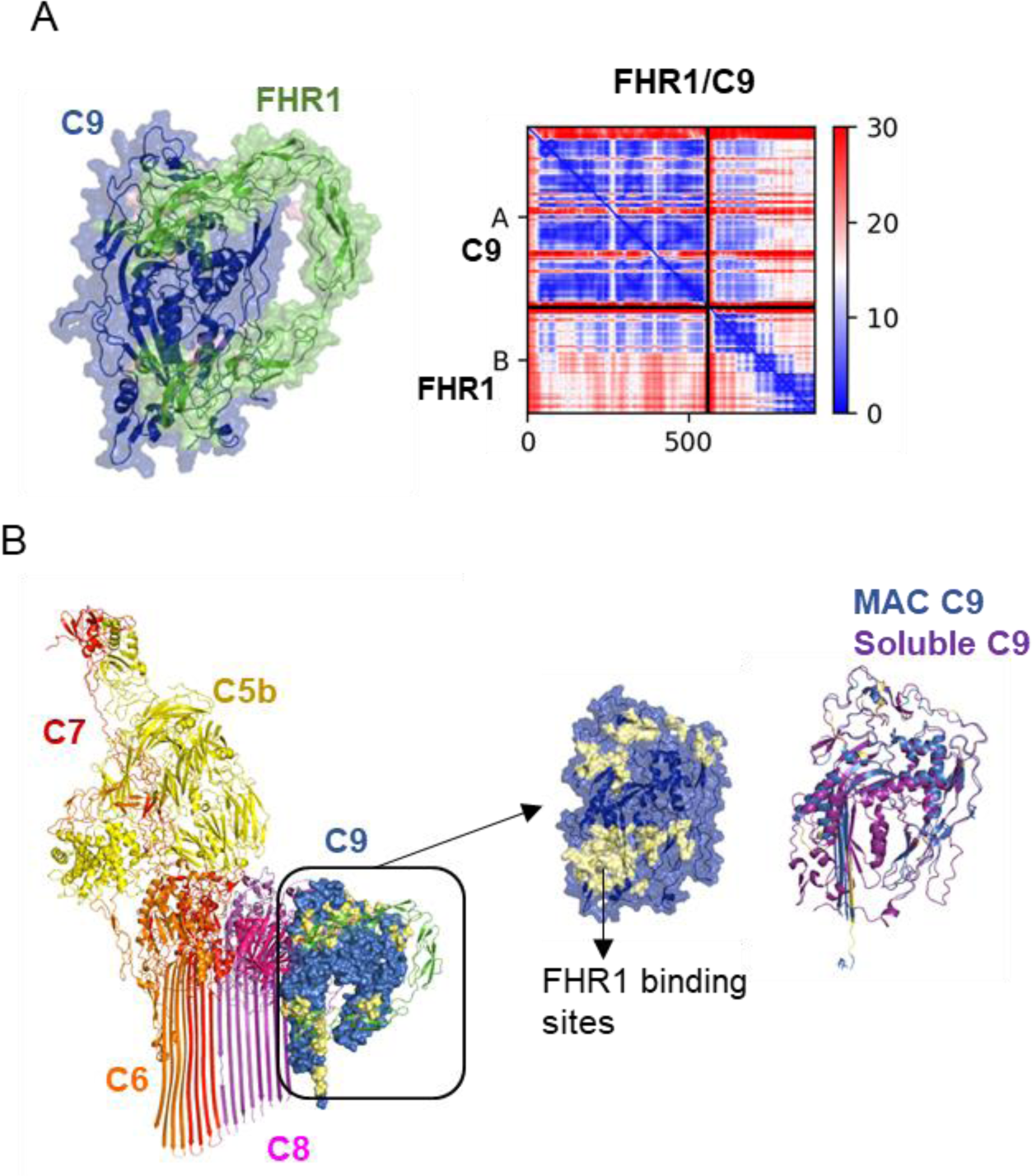
Interactions between C9 and FHR1 predicted by AFM-v1. **A.** Best-ranked model of FHR1 (green) in complex with C9 (blue) with interfaces located in FHR1_1-2_ and FHR1_4-5_. The binding interfaces predicted by AFM-v1 involved 13, 8, 7 and 12 residues in FHR1_1_, FHR1_2_, FHR1_4_ and FHR1_5_, respectively. On the right the Inter-chain Predicted Aligned Error (PAE) plot (20 recycles) with pLDDT = 83.6, ptmscore = 0.736 and iptm = 0.598. **B.** Superimposition of soluble C9 (purple) from model shown in A with C9 MAC (PDB 7nyc, dark blue) indicates that binding interfaces of FHR1 with C9 coincide with C9-C9 polymerization interface. FHR1 binding sites are shown in pale yellow. The FHR1_1-2_ binding interface forms 10 hydrogen bonds, and 8 hydrophobic interactions, while the FHR1_4-5_ forms 9 electrostatic interactions, 16 hydrogen bonds and 4 hydrophobic interactions. The key residues in the top-ranked model predicted by AFM-v1 and identified by MM-GBSA implemented in the HawkDock server were Arg 291, Phe 66, Tyr 57, Trp 67, Ile 32, Lys 287, Asn 239, Pro 223, Arg 270, Phe 91, and Phe 88 in FHR1.

**Supplementary Table 1.** Quality scores of models predicted with AlphaFold-Multimer version 1 (AFM-v1) or version 2 (AFM-v2) with different recycle numbers.

|  | complex | Recycles | pLDDT | ptmscore | iptm |
| --- | --- | --- | --- | --- | --- |
| AFM-v1 |  |  |  |  |  |
|  | C9/FHR1 | 3 | 82.4 | 0.695 | 0.549 |
|  | C9/FHR1 | 6 | 82.8 | 0.725 | 0.587 |
|  | C9/FHR1 | 12 | 83.3 | 0.727 | 0.602 |
|  | C9/FHR1 | 20 | 83.6 | 0.736 | 0.598 |
| AFM-v2 |  |  |  |  |  |
|  | C9/FHR1 | 3 | 74 | 0.579 | 0.272 |
|  | C9/FHR1 | 6 | 74.4 | 0.579 | 0.248 |
|  | C9/FHR1 | 12 | 72.4 | 0.573 | 0.196 |
|  | C9/FHR1 | 20 | 73.9 | 0.577 | 0.24 |
| AFM-v1 |  |  |  |  |  |
|  | C7/FHR1 | 3 | 80.9 | 0.618 | 0.446 |
|  | C7/FHR1 | 6 | 83.1 | 0.716 | 0.651 |
|  | C7/FHR1 | 12 | 82.8 | 0.733 | 0.668 |
|  | C7/FHR1 | 20 | 82 | 0.699 | 0.516 |
| AFM-v2 |  |  |  |  |  |
|  | C7/FHR1 | 3 | 70.7 | 0.5 | 0.355 |
|  | C7/FHR1 | 6 | 74.6 | 0.52 | 0.226 |
|  | C7/FHR1 | 12 | 74.5 | 0.518 | 0.2 |
|  | C7/FHR1 | 20 | 72.2 | 0.512 | 0.303 |
| AFM-v1 |  |  |  |  |  |
|  | C3d/FHR1 | 3 | 90.5 | 0.829 | 0.897 |
|  | C3d/FHR1 | 6 | 90.5 | 0.829 | 0.897 |
|  | C3d/FHR1 | 12 | 90.7 | 0.843 | 0.899 |
|  | C3d/FHR1 | 20 | 89.9 | 0.826 | 0.89 |
| AFM-v2 |  |  |  |  |  |
|  | C3d/FHR1 | 3 | 85.4 | 0.726 | 0.866 |
|  | C3d/FHR1 | 6 | 86 | 0.744 | 0.868 |
|  | C3d/FHR1 | 12 | 86 | 0.751 | 0.87 |
|  | C3d/FHR1 | 20 | 86 | 0.757 | 0.87 |

## Notes

### Competing Interest Statement

The authors have declared no competing interest.

